# Octopamine regulates neural circuits in the mushroom body and central complex, influencing sleep and context-dependent arousal

**DOI:** 10.1101/2025.03.06.641836

**Authors:** Martin Reyes, Yi Shen Lee, Preeti Sundaramurthi, Namrata Dhungana, Amanda Nguyen, Divya Sitaraman

**Affiliations:** Department of Psychology, College of Science, California State University- East Bay, Hayward, CA United States

## Abstract

Sleep is a complex and ubiquitous phenomenon in the animal kingdom, yet the mechanisms, modes, and effects of sleep on the brain and body remain highly variable and not fully understood. While sleep and wakefulness are often measured as two binary states at the behavioral level, animals exhibit a wide range of arousal states during wakefulness. Behaviors such as feeding, courting, escaping predators, or avoiding unfavorable environments are critical for survival and often compete with sleep. Biogenic amines, including dopamine, norepinephrine, and serotonin, are known to regulate sleep and arousal behaviors across species, offering valuable insights into how these states are co-regulated.

In this study, we leverage the small number, discrete organization, and well-defined connectivity patterns of neuromodulatory neurons in the fly brain to investigate how specific octopamine (OA) neurons regulate sleep and context-dependent arousal. We focus on a pair of OA neurons, VPM3, which project extensively to the mushroom body (MB) and central complex (CX). Our findings reveal that VPM3 neurons are sexually monomorphic and that their activity is essential for sleep suppression and male courtship behavior. Furthermore, we demonstrate that the male-specific form of the fruitless gene in these neurons plays a critical role in sleep regulation and that their activity is influenced by sleep history. Using connectome data and highly specific genetic tools, we identify upstream inputs to VPM3 neurons from the CX and elucidate the downstream pathways mediated by the MB, as well as the role of specific OA receptors. These detailed investigations highlight the multifaceted role of octopamine in sleep suppression and context-specific arousal, providing a crucial link between identified sleep microcircuits in the mushroom body and central complex.

## Introduction

Sleep is a neurobehavioral state that is highly conserved across species and required to maintain physiological and behavioral processes. As a quiescent state sleep prevents animals from engaging in goal directed behaviors like foraging, mating, escaping predators etc. that are critical for the survival of the organisms and the species. Hence, conflict between increasing wake hours and activity and need for sleep is continually encoded within the nervous system which in turn allows for these states to persist or break(reviewed in Beckwith and French, 2019; Eban-Rothschild, 2022; Eban-Rothschild et al., 2018, 2017; Saper et al., 2010, 2005; Sulaman et al., 2023).

The underlying neural mechanisms and interaction between systems that drive sleep, wakefulness and arousal are not well understood in any system. In both flies and mammals, neurons that regulate sleep and wakefulness are distributed throughout the brain (and ventral nerve cord, vnc in flies) and multiple approaches are beginning to illuminate how these neurons interact and integrate information to generate behavioral states of activity and quiescence(Joiner, 2018; Jones et al., 2023; Ni et al., 2019; Satterfield et al., 2022; Shafer and Keene, 2021).

Key biogenic amines like dopamine, serotonin, nor-epinephrine/octopamine, and histamine modulate multiple biological functions including learning and memory, circadian rhythms, aggression, mating, hunger, thermoregulation, and sleep. In *Drosophila*, a small number of cells (20-500) produce these aminergic modulators and innervate several regions of the fly brain and vnc. The extensive innervation patterns of these neurons uniquely position these molecules to influence multiple brain regions and enact state dependent behaviors in an efficient and robust way (Beckwith and French, 2019; Eban-Rothschild et al., 2018, 2017; Griffith, 2013; Helfrich-Förster, 2018; Jones, 2020; Liu and Dan, 2019; Saper et al., 2010).

The prevalent monoamine neurotransmitters in invertebrates studied in the context of sleep and other co-regulated behaviors are 5-HT (serotonin), DA (dopamine), OA (octopamine), and TA (tyramine). Dopamine and Octopamine have been shown to promote arousal and suppress sleep and exert their function by interacting with multiple circuits in the brain. Specifically, identified classes of PPL1 cluster of dopamine neurons project to and inhibit sleep promoting populations within the fan shaped body (Liu et al., 2012; Pimentel et al., 2016; Ueno et al., 2012). Subsets of PAM cluster project to Kenyon cells and MBONs to regulate anatomically and functionally well-defined microcircuits to regulate wakefulness(Driscoll et al., 2021; Nall et al., 2016; Sitaraman et al., 2015b). Like dopamine, octopamine is also shown to increase arousal and decrease sleep but only a few specific OA mediated microcircuits have been studied in the context of sleep. Octopaminergic ASM neurons project to the insulin like peptide producing pars intercerebralis (PI) and require OAMB receptors (Crocker et al., 2010).

These PI neurons also show altered potassium channel function and increase in cAMP in response to octopamine. However, the sleep suppression induced by ASM activation is much smaller than broader activation effects of all octopamine neurons and other octopamine receptors have also been implicated in sleep regulation (Crocker et al., 2010; Crocker and Sehgal, 2008). More recently a small number of octopaminergic neurons, named MS1 (Male Specific 1), were shown to regulate the decision between sleep and courtship in males. Specifically activating MS1 neurons reduced sleep in males, and silencing MS1 neurons led to decreased female-induced sleep loss and impaired mating behavior (Machado et al., 2017).

We specifically focus on a pair of OA neurons, VPM3 that have extensive projections in the MB and CX. We find that VPM3 neurons are a class of sexually monomorphic neurons, and the activity of these neurons is required for sleep-suppression and male courtship. Further we show that manipulation of male form of fruitless in these neurons is critical to the sleep function and their activity is affected by sleep history. Using connectome data and highly specific targeting tools we pinpoint the upstream signals to these neurons from the CX and comprehensively elucidate the MB dependent downstream pathways and role of specific OA receptors. These detailed investigations of the OA-VPM3 neurons provide a key link between identified sleep microcircuits in MB and CX.

## Results

### OA VPM3 neurons promote wakefulness

The mushroom body (MB) contains approximately 2,000 Kenyon cells (KCs), which are divided into seven anatomical classes based on the specific projections of their axons to the α/β, α′/β′, or γ lobes. These lobes are innervated by distinct classes of inputs, most notably dopamine neurons, anterior paired lateral (APL) neurons, and dorsal paired medial (DPM) neurons. The dendrites of mushroom body output neurons tile the lobe compartments, and specific populations have been identified and characterized in sleep regulation (Aso et al., 2014b; Haynes et al., 2015; Joiner et al., 2006; Pitman et al., 2006; Sitaraman et al., 2015a). Dopaminergic neurons, including PAM and PPL neurons, project to specific sleep- and wake-promoting compartments of the mushroom body. These neurons play a critical role in state switching and the persistence of wake states (Driscoll et al., 2021; Nall and Sehgal, 2014; Nall et al., 2016; Sitaraman et al., 2015b; Titos et al., 2023).

In addition to dopamine, subsets of octopaminergic (OA) neurons also extensively innervate the mushroom body (MB), though their role in sleep regulation remains unexplored. Ventrally located OA neurons, OA-VPM3 and OA-VPM4 neurons broadly project to the MB and form extensive synapses with Kenyon cells (KCs), dopaminergic neurons (DANs), and MB output neurons (MBONs) key regions implicated in sleep regulation. Additionally, OA-VUMa2 neurons synapse onto KCs in the MB calyx (Busch et al., 2009; Selcho, 2024).

A potential role for OA neurons projecting to MB in modulating sleep is further supported by the finding that broader manipulation of octopamine neurons suppresses sleep. To elucidate the role of OA VPM 3 and 4 neurons which are largely pre-synaptic to MB we assayed multiple GAL4 and split-GAL4 lines that target both VPM3 and 4 (R24E06, MB022B), VPM3 (SS46630) or VPM4 (R95A10, MB021B, SS46603). We used UAS-dTRPA1 to conditionally activate neurons and found significant reduction in nighttime sleep in flies on day 2 (activation, 29 °C) as compared to baseline (day 1 at 21°C) (Figure 1A, D-F). Even though these neurons were activated for a 24-hour duration including daytime, the sleep suppression was specific to nighttime sleep (Figure 1G and H). In these and subsequent behavioral experiments, we used empty split-GAL4 w [1118]; P {p65-AD.Uw} attP40; P{GAL4-DBD.Uw} attP2 as controls which has enhancer less fragments inserted in the same chromosomal location as the split-GAL4s or GAL4s.

**Figure 1:**
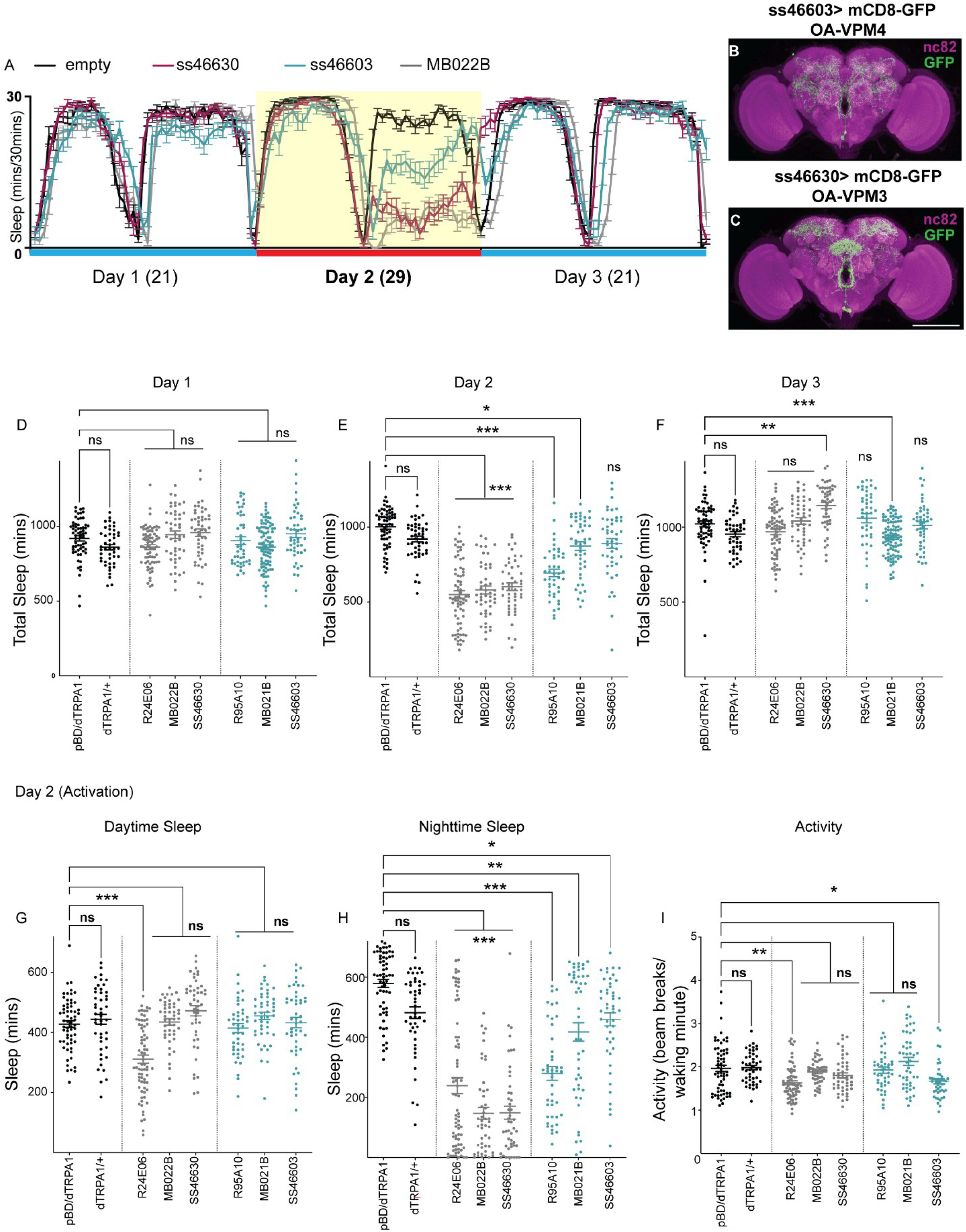
Activation of OA-VPM neurons promotes wakefulness and arousal. A. Sleep profiles of ss46630 (VPM3, red)>UAS-dTRPA1, ss46603 (VPM4, blue)> UAS- dTRPA1, MB022B (VPM3 and 4, grey)>UAS-dTRPA1 and pBD (empty, black)-split> UAS- dTRPA1. Sleep time plotted in 30 min bins and profile represents 3 days, 12h light and 12 h dark condition (day 1: 21, day 2: 29 (activation) and day 3: 21). B and C. Whole-mount brain immunostaining of ss46603 (VPM4)>UAS-mCD8-GFP and ss 46630 (VPM3)> UAS-mCD8-GFP flies with anti-GFP (green) and anti-Bruchpilot (BRP, nc82, magenta) antibody staining. Maximal intensity projection of the central brain is shown. Scale bars, 100 μm. D-F: Sleep duration on days 1, 2 and 3 of flies expressing dTRPA1 in broad and specific VPM3 drivers: ss46630 (n=47), MB022B (n=48), R24E06 (n=72), VPM4 drivers: ss46603 (n=46), MB021B (n=47), R95A10 (n=44) and genotypic controls (empty-split>UAS-dTrpA1 (n=62) and UAS-dTRPA1/+(n=47)). G, H: Sleep duration during daytime and nighttime on day 2 (activation) of the tested genotypes. I. Activity or number of beam counts/waking minute on day 2 (activation) is shown for the tested genotypes. For figures D-I, Mean ± SEM is shown and comparisons are made using Kruskal-Wallis test followed by Dunns multiple comparisons test. Statistical significances are indicated as ∗p < 0.05; ∗∗p < 0.01; ∗∗∗p < 0.001; ns, not significant. Figures A, D-I show male flies.

**Figure 1_Supplement 1:**
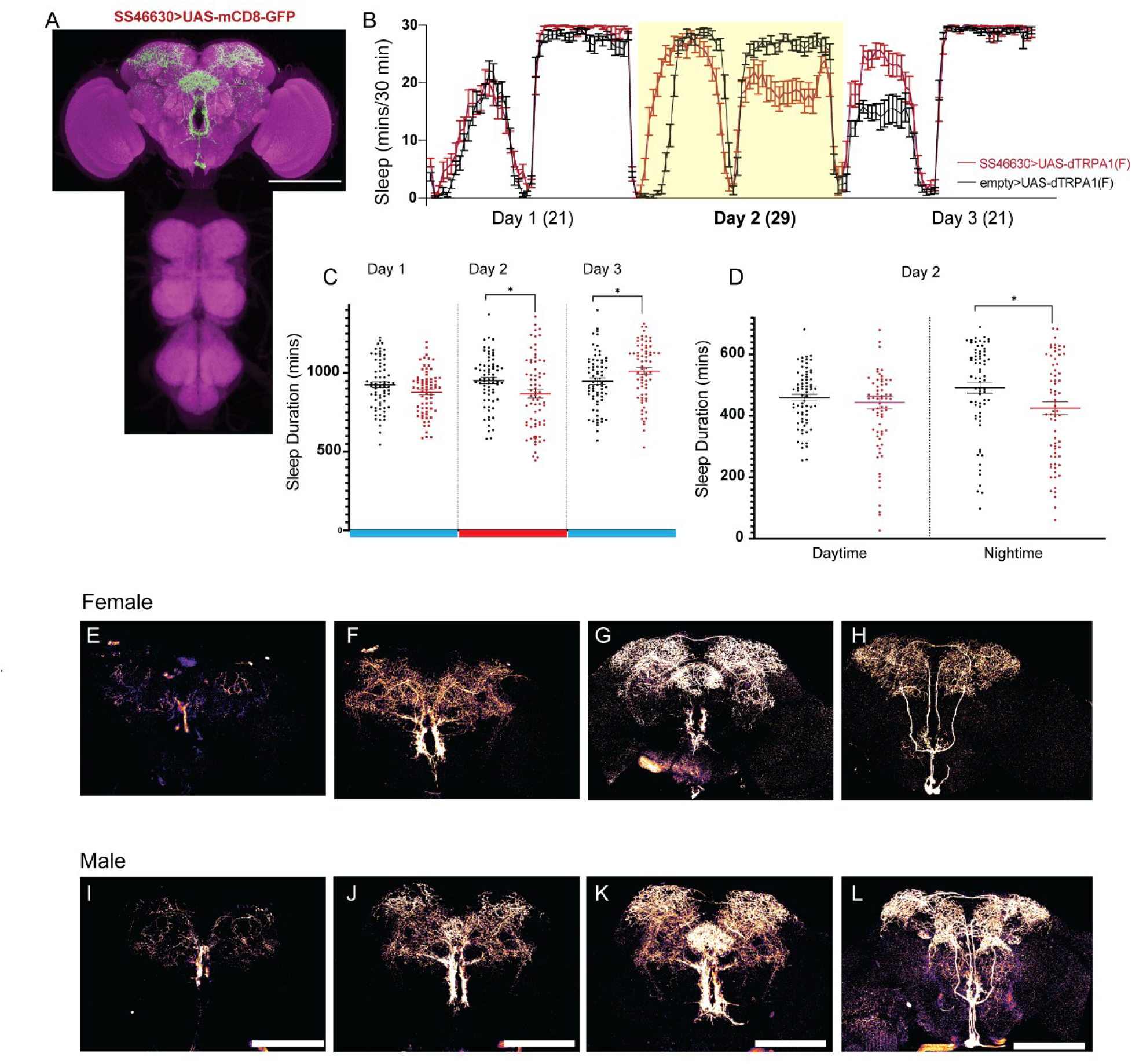
Arousal effects of OA-VPM3 activation are variable between male and female flies. A. Whole-mount brain and ventral nerve cord immunostaining of SS46630 (VPM3)>UAS-mCD8- GFP female fly. Maximal intensity projection of the central brain and vnc is shown. Scale bars, 100 μm. B. Sleep profiles of ss46630 (VPM3, red)>UAS-dTRPA1 and pBD (empty, black)-split> UAS- dTRPA1. Sleep time of female flies are plotted in 30 min bins and profile represents 3 days, 12h light and 12 h dark condition (day 1: 21, day 2: 29 (activation) and day 3: 21). C. Total sleep duration on days 1, 2 and 3 of female flies ss46630>UAS-dTRPA1 (n=70) and control empty-split>UAS-dTrpA1 (n=70). D. Sleep duration during daytime and nighttime on day 2 (activation) of the tested genotypes ss46630>UAS-dTRPA1 and empty-split>UAS-dTrpA1. For figures C and D, Mean ± SEM is shown and comparisons are made using Mann-Whitney U test. Statistical significances are indicated as ∗p < 0.05. Figures A-D show female flies. **E-H:** Whole-mount brain immunostaining of ss46630 (VPM3)> UAS-mCD8-GFP female flies with anti-GFP (green) stacks represented from posterior to anterior. Each image is maximal intensity projection of 15-20 (1um slices) pseudo colored to show the extensive innervation patterns within calyx, mushroom body, LH and SLP/SMP. E-H represent posterior to anterior slices **I-L:** Whole-mount brain immunostaining of ss46630 (VPM3)> UAS-mCD8-GFP male flies with anti-GFP (green) stacks represented from posterior to anterior. Each image is maximal intensity projection of 15-20 (1um slices) pseudocolored to show the extensive innervation patterns within calyx, mushroom body and SLP/SMP. I-L represent posterior to anterior slices Scale bar represents 100um.

To parse out the role of VPM3 and 4 neurons we tested driver lines specific to these subsets and found that the activation of VPM4 only had a modest effect on nighttime sleep. Since VPM4 lines R95A10 and MB021B have expression in VNC neurons in addition to labelling central brain VPM4 neurons it is unclear if the VNC neurons contribute to the phenotype. To address this, we generated a third specific driver for VPM4 (SS46603, Figure 1B) which has no VNC expression and only has a mild phenotype when activated with UAS-dTRPA1.

To test a more specific role for VPM 3 we generated an additional split-GAL4 line SS46630 which has no VNC expression or VPM4 neuron expression (Figure 1C, supplemental Figure S1A). Activation of SS46630 with dTRPA1 showed a strong decline in nighttime sleep recapitulating the phenotype of R24E06 and MB022B. Activation of VPM3 (using specific driver SS46630) shows a strong sleep rebound on day 3 after sleep loss induced by activation on day 2 (Figure 1F). In addition to sleep amounts, we also quantified activity (beam breaks/ waking minute) and found that decreased sleep was not a result of altered activity.

Indeed R24E06 (VPM3 and 4 driver) and SS46603 (VPM4 split-GAL4) had reduced activity (beam breaks/waking minute) as compared to their genotypic controls. Taken together, our systematic analysis of MB projecting octopamine neurons VPM3 and VPM4 using multiple overlapping and specific drivers with no VNC expression indicates a strong role for OA-VPM3 in wakefulness and arousal with a specific and more salient role in nighttime sleep.

In addition to testing males, we also tested age matched (3-8 days) female flies expressing UAS-dTRPA1 in OA-VPM3 neurons. Unlike males, female flies only had a modest decrease on nighttime sleep and weak sleep rebound post-activation (Figure S1B-D). We systematically compared VPM3 driver (SS46630) expression pattern in males and females expressing UAS- mCD8-GFP and found it to be identical or indistinguishable at the light microscopy level (Figure 1-supplement 1).

OA-VPM3 neurons are a single pair of neurons that are Tdc2+ and project from the anterior ventrolateral esophagus (oes) to the SMP and SLP neuropil passing the margin of the antennal lobe (al)(Busch et al., 2009). In addition to their extensive connectivity within MB, they also innervate the superior protocerebrum extensively, and the fan-shaped body (fb) (Figure 1-supplement 1), specifically FB6 layers that are part of the dorsal fan shaped body projections implicated in encoding of sleep drive (Donlea, 2017; Donlea et al., 2014, 2011; Pimentel et al., 2016).

Sexually dimorphic sleep phenotypes are not uncommon as male and female flies have different sleep patterns and octopaminergic neurons including the VPM subsets have been implicated in sex specific behaviors like courtship and aggression(Andrews et al., 2014; Certel et al., 2010; Manjila et al., 2019; Sherer et al., 2020). Further, VPM subsets are thought to express male isoform of fruitless (FruM) which might be the basis of sex specific behavioral modulation by OA neurons(Certel et al., 2010).

We next asked whether OA-VPM3 neurons are required for daily sleep. To address this, we expressed UAS-Kir2.1 in VPM3 neurons in male (Figure 2A-D) and female flies (Figure 2E-H) and compared sleep amount with genotypic controls. In addition to sleep amount (minutes/24- hour period, Figure 2A and 2E) we also looked at metrics of sleep consolidation, P(doze) (Figure 2B, F) and P(wake) (Figure 2C, G). P(wake) is the probability of transitioning from an inactive to an active state, whereas P(doze) is the probability of transitioning from active to inactive state(Wiggin et al., 2020). In males and females, inhibition of VPM3 increased sleep duration and decreased P(wake). However, P(doze) was not altered by inhibition in males but was reduced in females. Together these findings suggest that VPM3 neurons are wake-promoting and in addition to sleep duration alter sleep depth and pressure.

**Figure 2:**
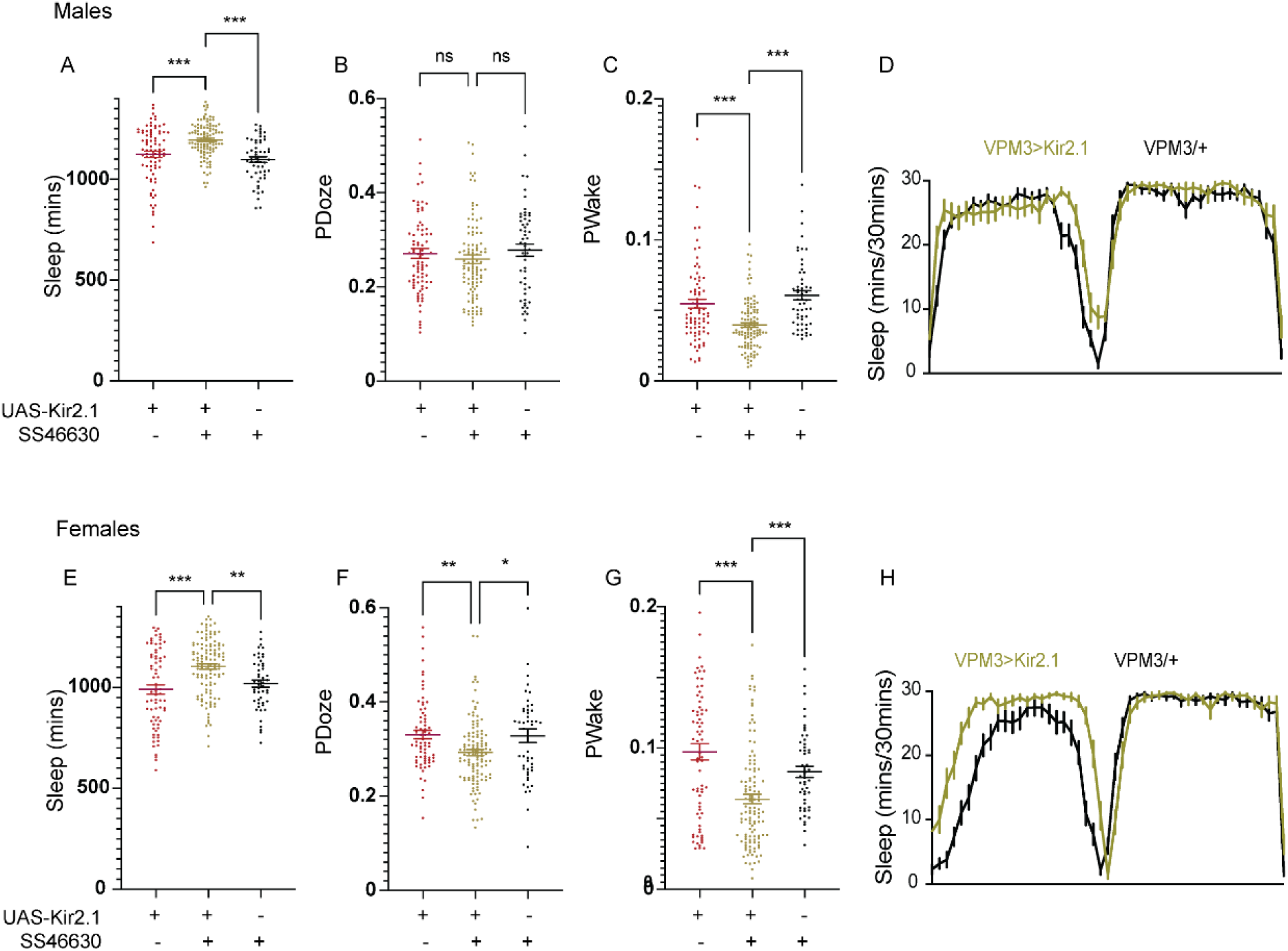
Inhibition of OA-VPM3 neurons increases sleep. A-C: Sleep duration, PDoze and PWake of ss46630>UAS-Kir2.1 (n=108) and controls (UAS- Kir2.1/+ (82) and ss46630/+(55)) in male flies. D: Sleep profiles of ss46630>UAS-Kir2.1 (green) and ss46630/+ (black) of male flies E-G: Sleep duration, PDoze and PWake of ss46630>UAS-Kir2.1 (n=118) and controls (UAS- Kir2.1/+ (72) and ss46630/+(51)) in female flies. H: Sleep profiles of ss46630>UAS-Kir2.1 (green) and ss46630/+ (black) of female flies In figures A-C and E-G, Mean ± SEM is shown and comparisons are made using Kruskal-Wallis test followed by Dunns multiple comparisons test. Statistical significances are indicated as ∗p < 0.05; ∗∗p < 0.01; ∗∗∗p < 0.001; ns, not significant.

### VPM3 neurons mediate context specific arousal

Numerous studies have addressed the role of octopamine neurons in motivated and goal directed sex-specific behaviors such as courtship and aggression(Certel et al., 2010; Dierick, 2008; Hoyer et al., 2008; Jia et al., 2021; Machado et al., 2017; Rohrscheib et al., 2015; Sherer et al., 2020; Stevenson et al., 2005; Watanabe et al., 2017; Zhou et al., 2008). However, less is known about how these behaviors are linked to sleep-suppression and how wakefulness is co-regulated with arousal and successful engagement in these motivated behaviors. Specifically, it’s unclear if distinct OA subsets regulate sleep-suppression and goal-directed arousal.

Sleep assays are conducted in single flies, making it difficult to directly observe how sleep deficits induced by octopamine activation influence other motivated behaviors. As a first step to understanding if VPM3 neuronal activation regulates other behaviors, we mined the publicly available dataset based on computer-vision-based unbiased quantification of 2,204 GAL4 lines from the Janelia GAL4 collection which uses the fly bowl assay (https://kristinbranson.github.io/BABAM/)(Robie et al., 2017)

We specifically focused on R24E06 which specifically labels OA VPM 3 and 4 neurons and has sparse expression in the VNC. Furthermore, activation of R24E06 with UAS-dTRPA1 suppresses sleep (Figure 1). In the fly bwol assay, activation of R24E06 with UAS-dTRPA1 in male-female groups shows increases in male wing extension, attempted copulation, touching and chase behavior as compared to controls. Activation of these neurons produced strong courtship arousal behaviors in males, with only minor changes in females. (Robie et al., 2017)

To test a more specific role of OA-VPM3 neurons in male-female courtship we tested flies expressing dTRPA1 in SS46630 (VPM3 specific driver, no VNC expression) at 21 and 29°C and found significant increases in total courtship duration, following, wing extension and attempted copulation during the 30-minute recording (Figure 3 A-D). Since, R24E06 also targets VPM4 we tested flies expressing dTRPA1 in VPM4 neurons driven by SS46603 and found that VPM4 neurons do not regulate courtship (Figure 3_supplement 1).

**Figure 3:**
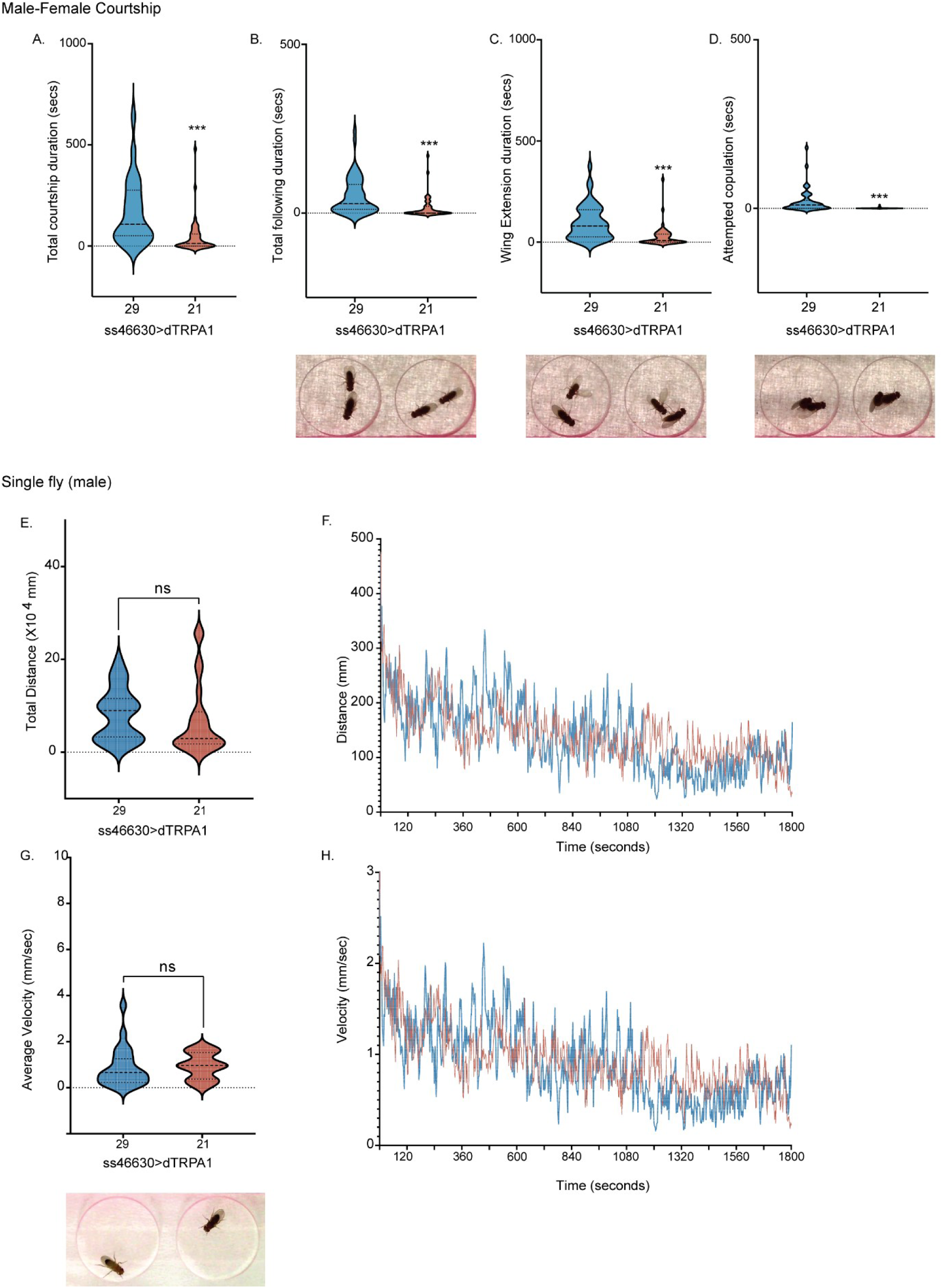
OA-VPM3 activation promotes context dependent arousal. A. Total courtship duration (SS46630>UAS-dTRPA1) during a 30-minute recording period at 21 (red) and 29 degrees (blue). n=48 flies, p<0.0001. B. Total following duration (SS46630>UAS-dTRPA1) during a 30-minute recording period at 21 (red) and 29 degrees (blue). n=48 flies, p<0.0001. sample image from recording showing following behavior. C. Total wing extension duration (SS46630>UAS-dTRPA1) during a 30-minute recording period at 21 (red) and 29 degrees (blue). n=48 flies, p<0.0001. sample image from recording showing wing extension behavior. D. Total attempted copulation duration (SS46630>UAS-dTRPA1) during a 30-minute recording period at 21 (red) and 29 degrees (blue). Two conditions were compared using unpaired t-test (Mann-Whitney U test) n=48 flies, p<0.0001. sample image from recording showing attempted copulation behavior. E. Total distance travelled by single flies (SS46630>UAS-dTRPA1) during a 30-minute recording period at 21 (red) and 29 degrees (blue). n=24 flies, p=0.7491. F. Average distance travelled during 30-minute recording period at 21 (red) and 29 degrees (blue). G. Average velocity of single flies (SS46630>UAS-dTRPA1) during a 30-minute recording period at 21 (red) and 29 degrees (blue). n=24 flies, p=0.5063. H. Average velocity during 30-minute recording period at 21 (red) and 29 degrees (blue). Violin plots represent the median and quartiles and two conditions were compared using unpaired t-test (Mann-Whitney U test).

**Figure 3_Supplement 1:**
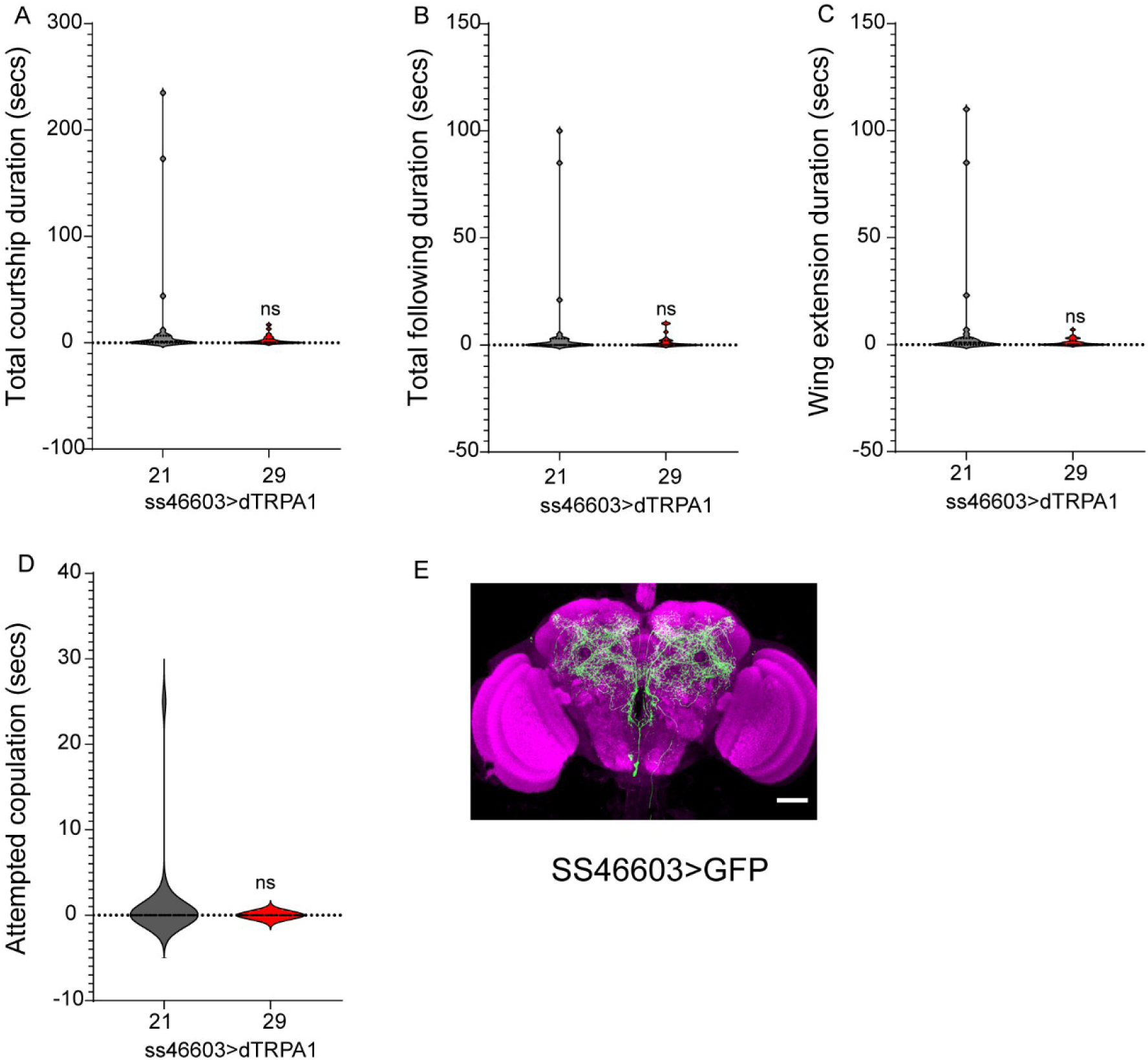
OA-VPM4 activation does not change male-female courtship behavior. A. Total courtship duration (SS46603>UAS-dTRPA1) during a 30-minute recording period at 21 (grey) and 29 degrees (red). n=24 flies. B. Total following duration (SS46603>UAS-dTRPA1) during a 30-minute recording period at 21 (grey) and 29 degrees (red). n=24 flies. C. Total wing extension duration (SS46603>UAS-dTRPA1) during a 30-minute recording period at 21 (red) and 29 degrees (blue). n=24 flies. D. Total attempted copulation duration (SS46603>UAS-dTRPA1) during a 30-minute recording period at 21 (red) and 29 degrees (blue). Two conditions were compared using unpaired t-test (Mann-Whitney U test) n=24 flies. E. Whole-mount brain immunostaining of SS46603 (VPM4)>UAS-mCD8-GFP male fly. Maximal intensity projection of the central brain and vnc is shown. Scale bars, 20 μm.

**Figure 3_Supplement 2:**
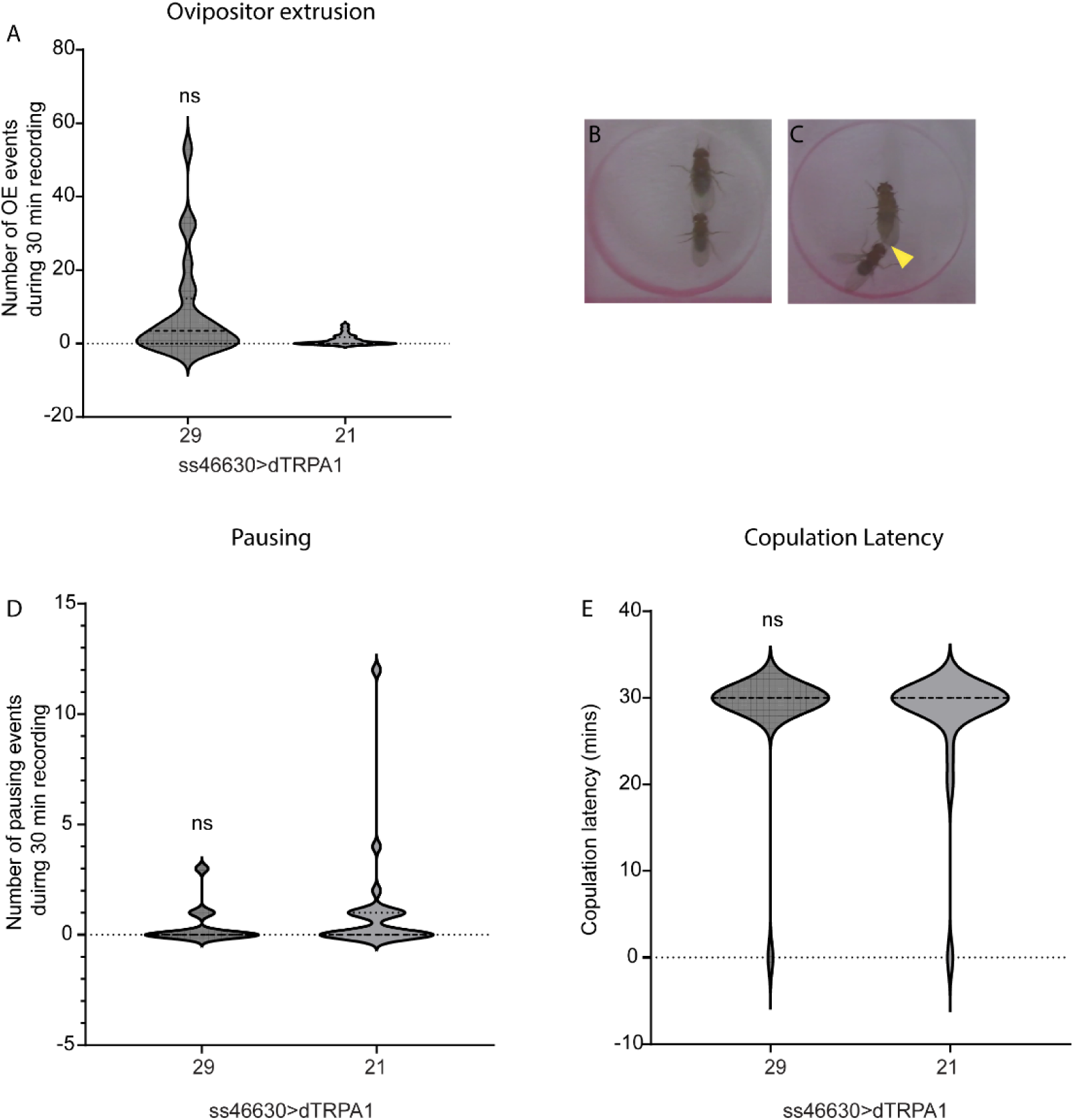
Female receptivity is not altered by OA-VPM3 activation. A. Number of ovipositor extrusion events during 30-minute recordings at 21 and 29 degrees. n=24 flies. SS46630>dTRPA1 female flies were tested with tester males. B and C. Sample image from recording showing pausing and ovipositor extrusion events C. Number of pausing events during 30-minute recording at 21 and 29 degrees. n=24 flies. D. Number of pausing events during 30-minute recordings at 21 and 29 degrees. n=24 flies.

We also tested SS46630>dTRPA1 male flies at 21 and 29°C in the same arena in the absence of female flies and did not find any changes in wing extension or other courtship parameters (Figure 3E-H). Additionally, we also measured locomotor parameters like total distance and average velocity during the 30-minute recording in the same arena used in male-female experiments described above. Activation of OA-VPM3 in male flies did not elicit the courtship specific behaviors or alter distance travelled and velocity as compared to controls (Figure 3E-H).

To test if VPM3 neurons are also involved in female courtship arousal, we tested SS46630>dTRPA1 female flies at 21 and 29°C with tester males and measured female receptivity behaviors (pausing, courtship latency and ovipositor extrusion(von Philipsborn et al., 2023; Yang et al., 2023b). Activation of VPM3 did not alter these behaviors and there were no significant differences in female receptivity behaviors at 21 and 29°C (Figure 3_supplement 2).

Taken together, we find that in addition to sleep-suppression, VPM3 neurons promote context specific courtship arousal in male but not female flies. Given the strong effects of VPM3 on sleep suppression and courtship arousal we focused on characterization of molecular regulators and output pathways in male flies.

### Sleep suppression by VPM3 is FruM dependent

The expression of male isoform of fruitless is thought to specify the neuronal circuitry required for male courtship behavior (Manoli et al., 2005; Stockinger et al., 2005), however it’s not clear if fruM expression in specific neurons is also involved in male sleep behavior. Like courtship, sleep behavior is sexually dimorphic but the differences in male and female sleep are subtle and depend on other variables like mating status, diet, age etc.

We recently investigated the role of FruM in male sleep and found that knockdown of FRUM in all fruM positive neurons targeted by fru-GAL4 significantly alters sleep duration(Chen et al., 2017). Additionally, several clusters of OA neurons targeted by Tdc2-Gal4 including VPM clusters are FruM positive and thought to controlsex specific motivated behaviors (Andrews et al., 2014; Certel et al., 2010; Sherer et al., 2020). Based on these data, we argued that fruM potentially functions within the OA-VPM3 neurons and regulates sleep differentially in male flies.

To address this directly, we co-expressed dTRPA1 and FruM-RNAi in VPM3 neurons and tested them at 21 and 29°C. We used two activation conditions lasting 24 hours (day and night) (Figure 4A-C) or 12 hours (Figure 4D-F) during nighttime as the effect of VPM3 activation was specific to nighttime sleep. We compared baseline and activation conditions between VPM3 and empty split-GAL4 flies expressing either UAS-dTRPA1, UAS-FruMRNAi or both UAS-dTRPA1 and FruMRNAi. We found that dTRPA1 induced activation of VPM3 suppressed sleep and this sleep loss was reversed by knockdown of fruM. The reversal of sleep suppression was partial in 24-hour dTRPA1 activation conditions and complete in 12-hour nighttime activation conditions.

**Figure 4:**
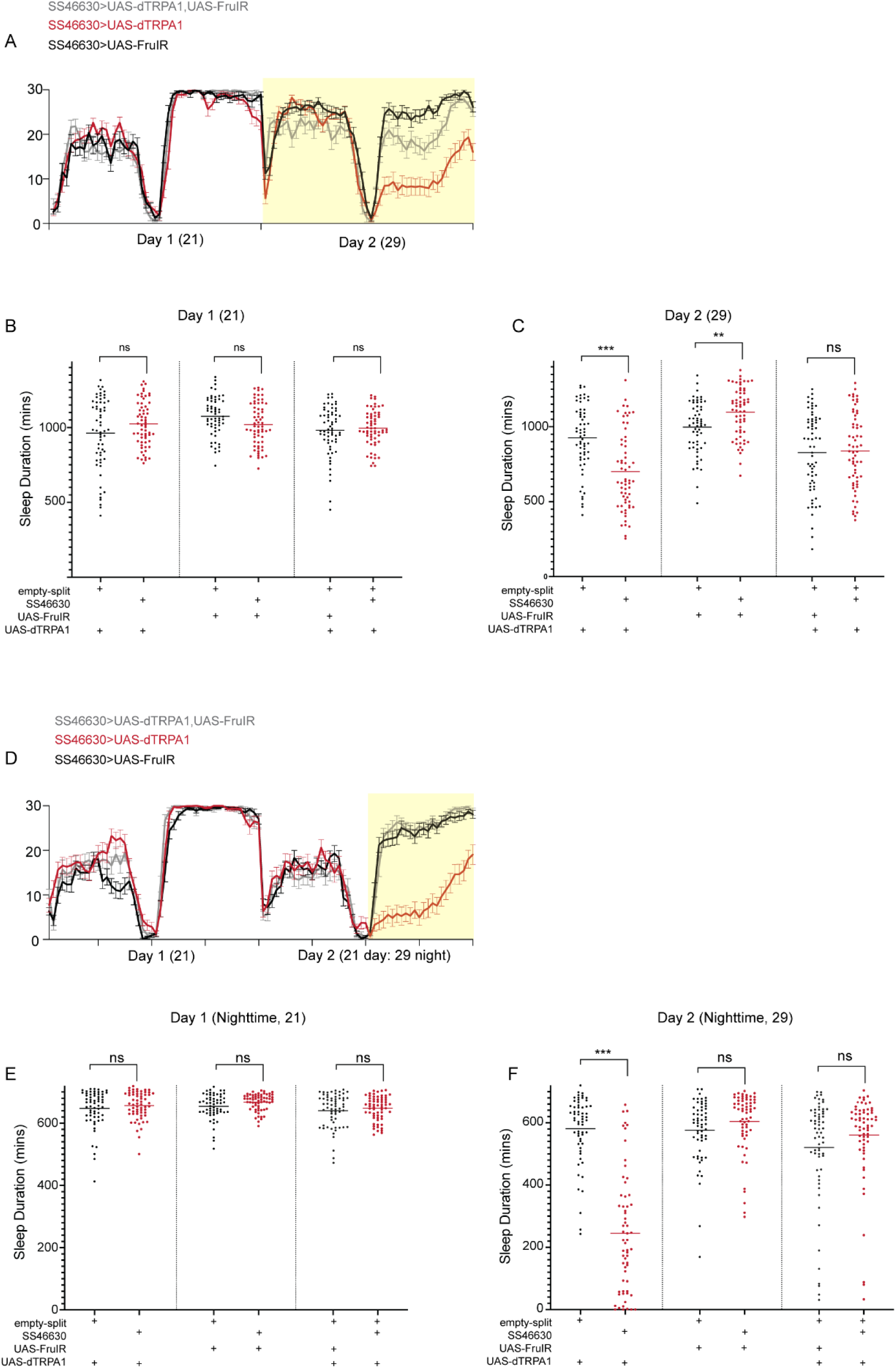
Sleep regulation by OA-VPM3 is fruM dependent. A. Sleep profile of flies (SS46630>dTRPA1, SS46630>FruMIR and SS46630>dTRPA1, FruMIR). Day 1 and Day 2 represent daytime and nighttime sleep at 21 and 29 degrees respectively. B. Sleep duration on day 1 (empty-split or SS46630 expressing UAS-dTRPA1, UAS-FruMIR or both) are plotted. C. Sleep duration on day 2 (empty-split or SS46630 expressing UAS-dTRPA1, UAS-FruMIR or both) are plotted. D. Sleep profile of flies (SS46630>dTRPA1, SS46630>FruMIR and SS46630>dTRPA1, FruMIR). Day 1 represents daytime and nighttime sleep at 21 and Day 2 represents daytime at 21 and nighttime at 29 degrees respectively. E. Nighttime sleep duration on day 1 (empty-split or SS46630 expressing UAS-dTRPA1, UAS- FruMIR or both) are plotted. F. Nighttime sleep duration on day 2 (empty-split or SS46630 expressing UAS-dTRPA1, UAS- FruMIR or both) are plotted. In figures B, C, E and F, Mean ± SEM is shown. Empty-split or SS46630 expressing identical transgenes were compared using Mann-Whitney U test. Statistical significances are indicated as ∗p < 0.05; ∗∗p < 0.01; ∗∗∗p < 0.001; ns, not significant. Genotypes used are: SS46630>dTRPA1 (n=62), SS46630>FruMIR (n=63), SS46630>dTRPA1, FruMIR (n=62), empty-split>dTRPA1 (n=61), empty-split>FruMIR (n=59) and empty-split>dTRPA1, FruMIR (n=63).

Flies expressing only FruM RNAi in VPM3 neurons showed mild increases in sleep during 24- hour temperature shift as compared to empty split-GAL4 controls, however we did not see this effect during 12-hour temperature shifts. These data show that FruM functions within VPM3 neurons to regulate sleep in male flies.

### Sleep need alters VPM3 activity in lateral and medial protocerebrum

Based on GFP expression and EM data, VPM3 neurons have significant inputs from central complex neurons innervating dorsal fan shaped body neurons, layer 6 and send significant outputs to the MB (KCs, DANs and MBONs), SMP and SLP region. VPM3 activation increases wakefulness and is followed by rebound sleep post-activation in males. Hence, VPM3 activation is sufficient to sleep deprive animals and induce subsequent homeostatic recovery sleep.

To directly test if sleep deprivation impacts the activity of VPM3 neurons we used CaLexA driven by 24E06-GAL4. The CaLexA system relies on a fluorescent reporter gene wherein calcium accumulation activates calcineurin that dephosphorylates NFAT, causing the chimeric transcription factor mLexA-VP16-NFAT to locate to the nucleus which in turn induces expression of GFP under the control of the LexAop (Masuyama et al., 2012).

Therefore, comparing the intensity of detectable fluorescence signal in sleep replete and sleep deprived flies can be used to understand how sleep needs modulate activity of VPM neurons. Sleep deprivation was induced by mechanical shaking for 12 hours (during nighttime) and to ensure that 24E06-GAL4>CaLexA flies were sleep deprived, we measured sleep during deprivation. Sleep replete flies were used as controls (Figure 5A and B). GFP fluorescence in whole mount brain was used to quantify difference in activity between two groups (sleep-deprived and rested flies) and we found significant differences in fluorescence intensity in the SMP/SLP region around the peduncle that are innervated by MBONs and DANs and dorsal layers of fan shaped body. Fluorescence in the ventral regions around the esophagus which have extensive VPM projections were highly variable between samples and were not significantly different between conditions.

**Figure 5:**
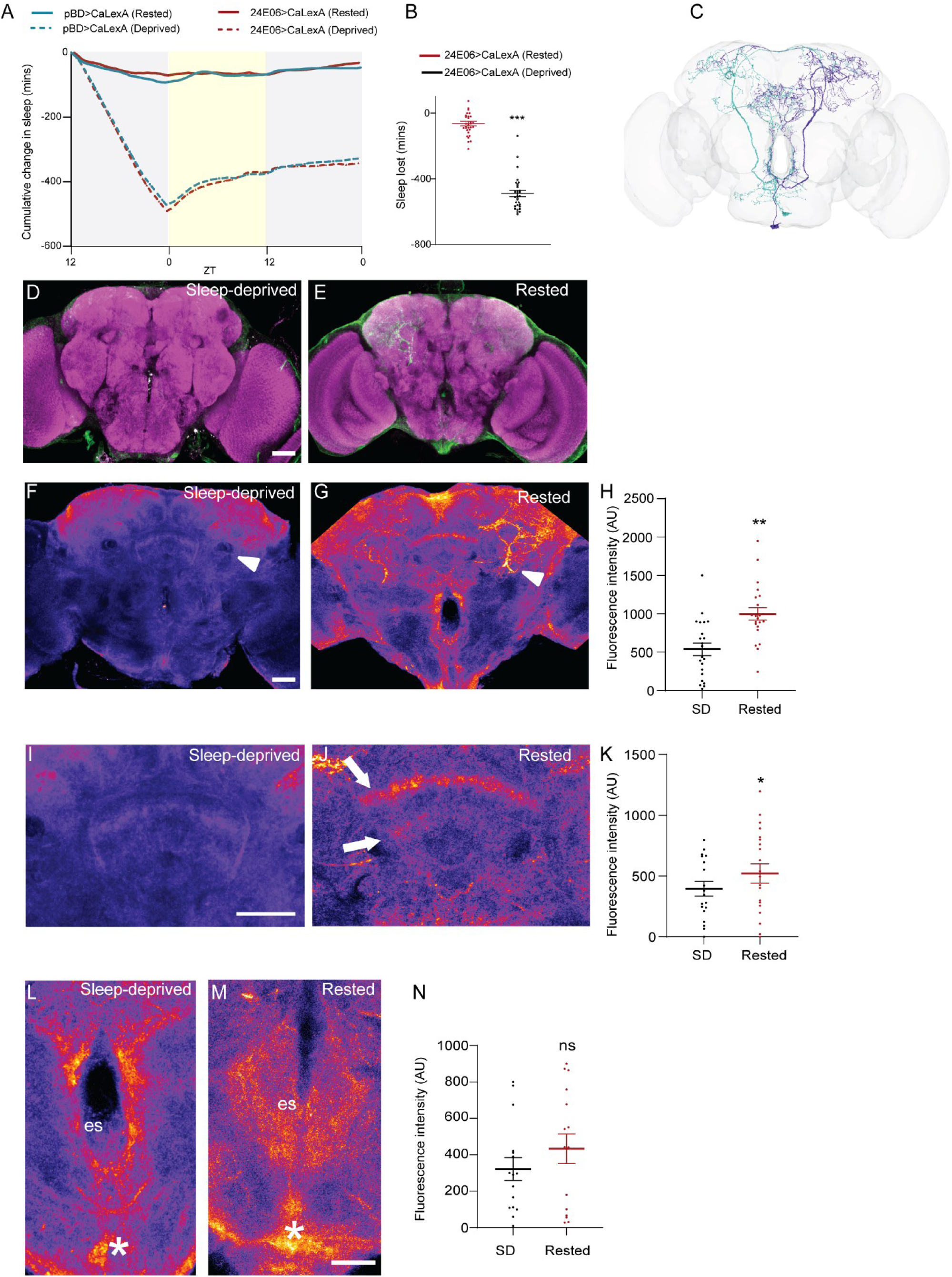
Sleep deprivation alters activity of VPM neurons in central brain regions. A. Cumulative sleep amount in minutes observed 12hr during deprivation and 24-hour recovery period (dashed line). Rested flies (solid lines) represent flies that were not deprived and monitored in parallel for the 36-hour duration. pBD and 24E06-Gal4 flies expressing CalexA are shown in blue and red respectively. B. Quantification of sleep lost (in minutes). Data represents 24E06-CalexA flies that were deprived (sleep deprived) and non-deprived controls (rested). Statistical comparisons are made by Mann-Whitney U test, statistical significances are indicated as ∗p < 0.05; ∗∗p < 0.01; ∗∗∗p < 0.001; ns, not significant. C. EM based skeleton reconstruction of OA-VPM3 neurons, projections and innervated brain regions (adapted from https://codex.flywire.ai/)(Dorkenwald et al., 2024; Schlegel et al., 2024) D and E. Whole-mount brain immunostaining of 24E06-GAL4>CalexA flies (sleep-replete controls (B), and sleep-deprived controls (C)) with anti-GFP (green) and anti-Bruchpilot (BRP, nc82, magenta) staining. Maximal intensity projection of the central brain is shown. Scale bars, 100 μm. F and G. Whole-mount brain immunostaining of 24E06-GAL4>CalexA flies (sleep-replete controls (E), and sleep-deprived controls (F)). GFP staining is pseudo colored for visualization. Maximal intensity projection of the central brain (10-15 slices, 1 um) is shown. Arrow indicates region around the peduncle to highlight SMP/SIP projections of OA-VPM3 neurons H. Quantification of CaLexA (fluorescence intensity) in the SMP/SIP region in flies (sleep-replete and sleep-deprived). Statistical comparisons are made by Mann-Whitney U test, statistical significances are indicated as ∗p < 0.05; ∗∗p < 0.01; ∗∗∗p < 0.001; ns, not significant. I and J. Whole-mount brain immunostaining of 24E06-GAL4>CalexA flies (sleep-replete controls (E), and sleep-deprived controls (F), central complex region are shown. GFP staining is pseudo colored for visualization. Maximal intensity projection of the central brain (8-10, 1 um) is shown. Arrows indicate dorsal and ventral projections of fan shaped body. K. Quantification of CaLexA (fluorescence intensity) in the central complex region in flies (sleep-replete and sleep-deprived). Statistical comparisons are made by Mann-Whitney U test, statistical significances are indicated as ∗p < 0.05; ∗∗p < 0.01; ∗∗∗p < 0.001; ns, not significant. L and M. Whole-mount brain immunostaining of 24E06-GAL4>CalexA flies (sleep-replete controls (E), and sleep-deprived controls (F)), central complex region is shown. GFP staining is pseudo colored for visualization. Maximal intensity projection of the central brain (8-10, 1 um) is shown. Asterisks indicate VPM cell bodies and es indicates esophagus. N. Quantification of CaLexA (fluorescence intensity) in the ventral region (sleep-replete and sleep-deprived). Statistical comparisons are made by Mann-Whitney U test, statistical significances are indicated as ∗p < 0.05; ∗∗p < 0.01; ∗∗∗p < 0.001; ns, not significant. Scale bars indicate 20um.

Since 24E06-GAL4 labels both VPM3 and VPM4 neurons, we tried using SS46630 to target CaLexA to VPM3 neurons. The GFP expression was weak to undetectable potentially because SS46630 is a weaker driver than 24E06-GAL4. VPM3 and VPM4 neurons have overlapping expression in SMP/SLP region and mushroom body lobes but projections in the fan shaped body are restricted to VPM 3 neurons. These data suggest that although we cannot completely exclude the role of VPM 4 in encoding sleep drive, sleep need alters VPM3 activity in key central brain regions.

### VPM3 neurons regulate sleep via interconnected MB input-output pathways

Based on light microscopy data VPM3 neurons dendrites extend within the sub- and peri-esophageal zones and project to the MB, CX and other brain regions(Busch et al., 2009; Sayin et al., 2019) (Figure 1_supplement 1). We probed the recently published EM volume (hemibrain, Neuprint) of the adult brain to identify specific connectivity between VPM3 and other brain regions(Zheng et al., 2018). We focused on inputs and outputs to and from the VPM3 neurons and sorted them based on number to synapses to identify other neurons that form strong connections with VPM3 (Supplementary Table 1, Figure 6A and B). We identified 2 MBONs, 1 class of Kenyon cells and several dopamine neuron clusters in the top 10 outputs. The strongest inputs to the VPM3 neurons based on the EM data are from several dorsal and ventral fan shaped body neurons.

**Figure 6:**
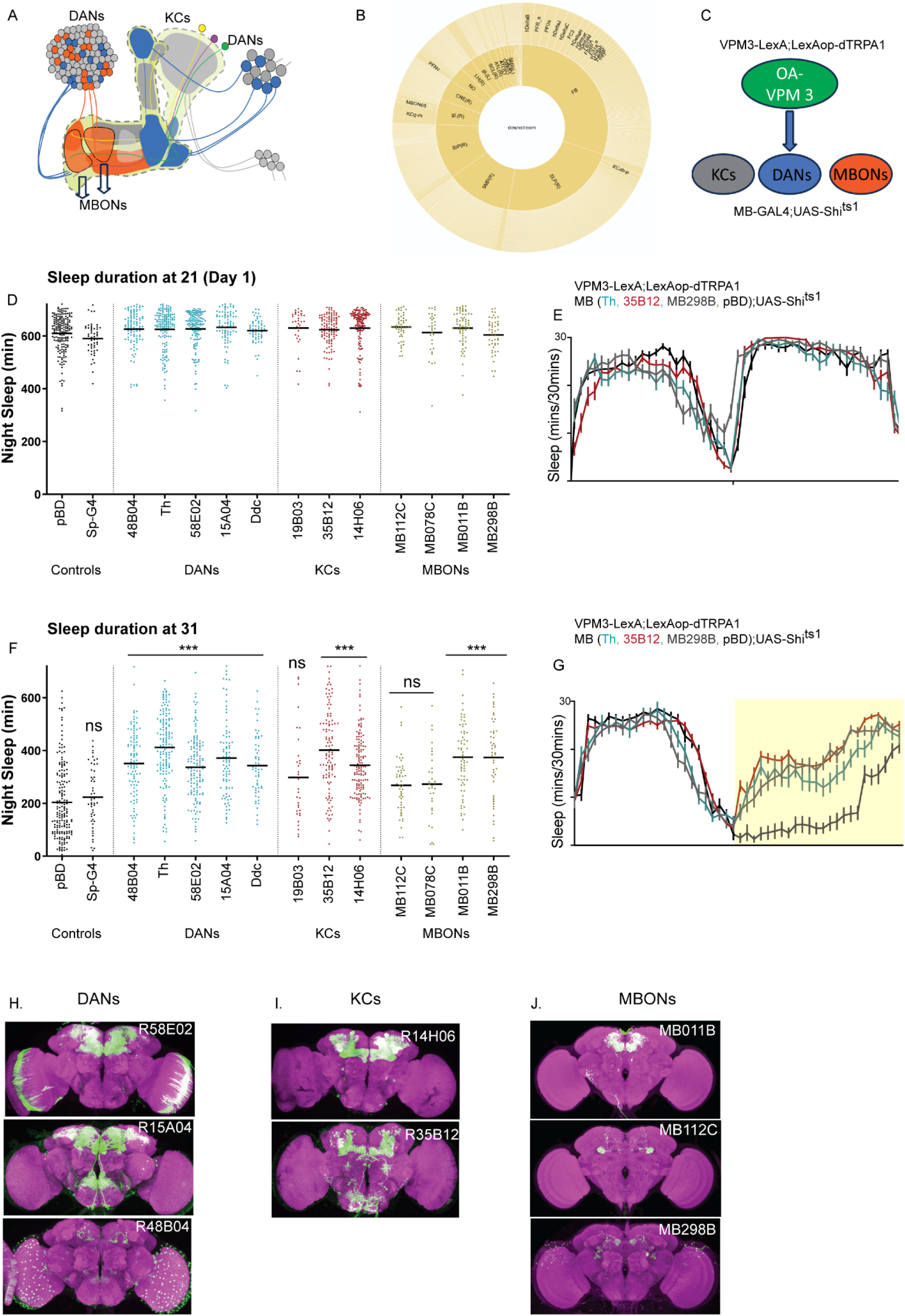
OA-VPM3 dependent arousal is mediated by sleep regulating neurons within mushroom body. A. Schematic of mushroom body Kenyon cell (KC) bodies are represented grey and blue circles depending on the projection lobe. KCs dendrites extend within the calyx and axons project to form α/β, α’/β’, and γ lobes. PAM and PPL1 dopaminergic neurons (DANs) extensively innervate the lobes. Mushroom body output neurons (MBONs) transmit signals from the mushroom bodies to various other brain regions. B. OA-VPM3 (hemibrain id: 329566174 and 5813061260) output represented as a starburst pattern from hemibrain connectome (NeuronBridge (janelia.org) shows extensive downstream signaling via γ lobes, PAM-DANs and specific MBONs (MBON11, MBON 5). C. Experimental strategy to study neuronal pairs by simultaneously activating VPM3 neurons (24E06-LexA; LexAop-dTRPA1) and silencing potential downstream neurons (X-Gal4; UAS- Shi^ts1^). D. Nighttime sleep duration (day 1, baseline at 21) of flies expressing 24E06-LexA; LexAop-dTRPA1 and X-Gal4; UAS-Shi^ts1^. X-Gal4 includes DANs (48B04 n=99,15A04 n=83, 58E02 n=134, TH n=147 and Ddc n=58), KCs (14H06 n=138, 19B03 n=32 and 35B12 n=109) and MBONs (MB078C n=32, MB112C n=50, MB298B n=48 and MB011B n=67). Controls include empty-split Gal4 (n=180) and w1118 (n=55). E. Sleep profile of selected lines showing daytime and nighttime sleep at 21. F. Nighttime sleep duration (day 2, 31) of flies expressing 24E06-LexA; LexAop-dTRPA1 and X- Gal4; UAS-Shi^ts1^. X-Gal4 includes DANs (48B04 n=99,15A04 n=83, 58E02 n=134, TH n=147 and Ddc n=58), KCs (14H06 n=138, 19B03 n=32 and 35B12 n=109) and MBONs (MB078C n=32, MB112C n=50, MB298B n=48 and MB011B n=67). Controls include empty-split Gal4 (n=180) and w1118 (n=55). G. Sleep profile of selected lines showing daytime (21) and nighttime sleep at 31. H-J. Whole-mount brain immunostaining of DANs (H), KCs (I) and MBONs (J) expressing 10X- UAS-mCD8-GFP flies with anti-GFP (green) and anti-Bruchpilot (BRP, nc82, magenta) antibody staining. Maximal intensity projection of the central brain was made from original z stack files obtained from https://flweb.janelia.org/cgi-bin. In figures D and F, mean is shown and comparisons are made using Kruskal-Wallis test followed by Dunns multiple comparisons test. Statistical significances are indicated as ∗p < 0.05; ∗∗p < 0.01; ∗∗∗p < 0.001; ns, not significant. For each genotype no data was excluded, and data collected represents 3 or more independent trials.

**Figure 6_Supplement 1:**
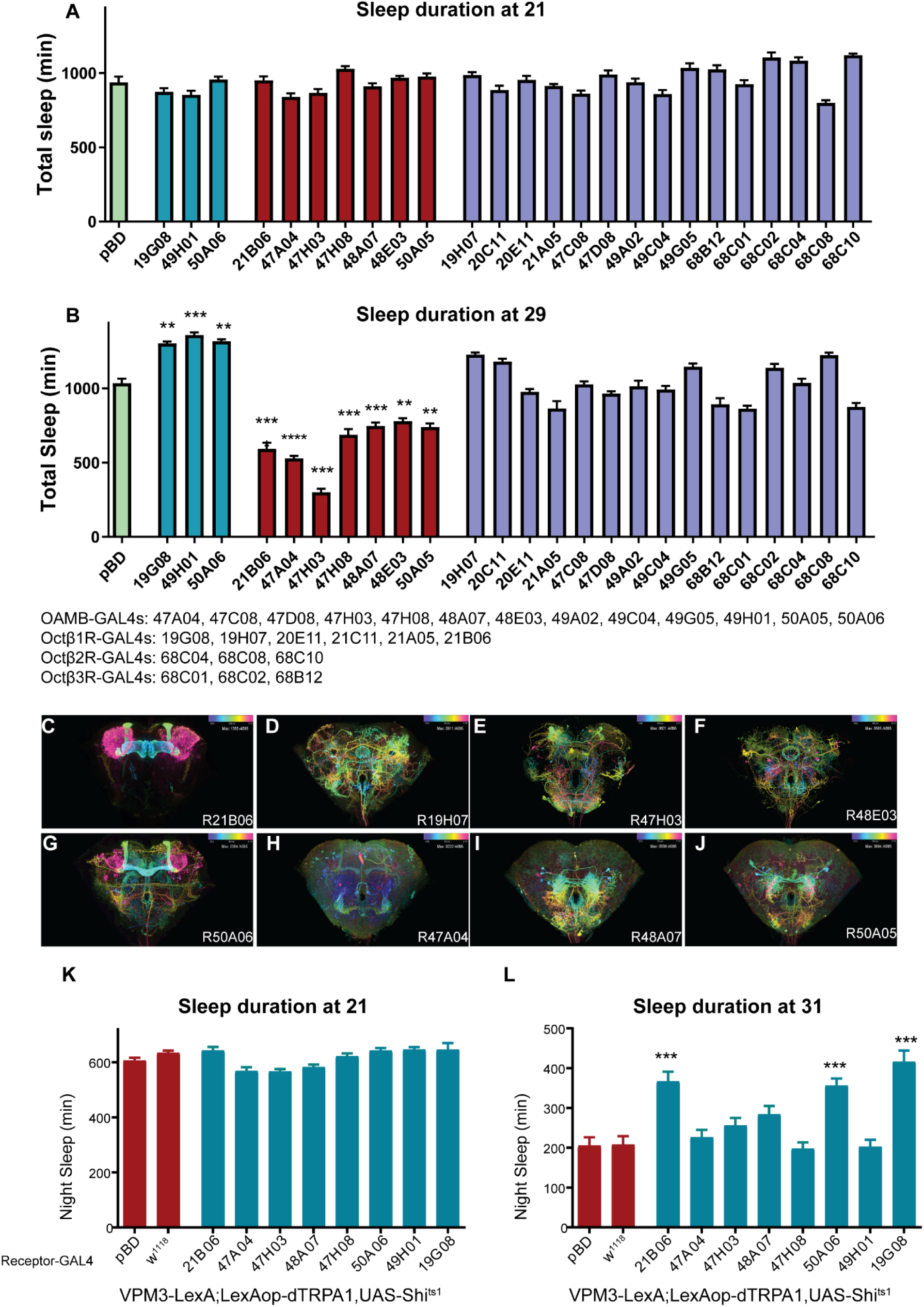
Behavioral and anatomical screening of OA Receptor-GAL4 lines for sleep phenotypes. A and B. Flies expressing UAS-dTRPA1 and 25 octopamine receptor GAL4 lines were tested at 21 and 29 degrees to identify receptor-GAL4 lines involved in sleep regulation.). Empty-GAL4 expressing dTRPA1 was used as a genotypic control and data is represented as mean and SEM and comparisons are made by Kruskal-Wallis test followed by Dunns multiple comparisons test. Statistical significances are indicated as ∗p < 0.05; ∗∗p < 0.01; ∗∗∗p < 0.001; ns, not significant. For each genotype no data was excluded, and data collected represents 2 or more independent trials. C-J. Whole-mount brain immunostaining of 8 octopamine receptor GAL4 lines with altered sleep on activation and sparse expression patterns. OA receptor GAL4 lines are expressing. Maximal intensity projection of the central brain was made from original z stack files obtained from https://flweb.janelia.org/cgi-bin. K. Nighttime sleep duration (day 1, baseline at 21) of flies expressing VPM3-LexA; LexAop-dTRPA1 and X-Gal4; UAS-Shi^ts1^. X-Gal4 represents 8 OA receptor GAL4 lines (R21B06, R47A04, R47H03, R48A07, R47H08, R50A06, R49H01 and R19G08). Controls include empty-Gal4 (pBD) and w1118. K. Nighttime sleep duration (day 2, 29 degrees)) of flies expressing VPM3-LexA; LexAop-dTRPA1 and X-Gal4; UAS-Shi^ts1^. X-Gal4 represents 8 OA receptor GAL4 lines (R21B06, R47A04, R47H03, R48A07, R47H08, R50A06, R49H01 and R19G08). Controls include empty-Gal4 and w1118. For K and L, and data is represented as mean and SEM and comparisons are made by Kruskal-Wallis test followed by Dunns multiple comparisons test. Statistical significances are indicated as ∗p < 0.05; ∗∗p < 0.01; ∗∗∗p < 0.001; ns, not significant. For each genotype no data was excluded, and data collected represents 2 or more independent trials.

Expression of UAS-syt-eGFP and UAS-Denmark in Tdc2-GAL4 neurons shows extensive input and outputs in fan shaped body neurons (Ni et al., 2019). To confirm whether the precise Tdc+ dfb outputs are indeed OA-VPM3 we conducted GRASP staining (GFP reconstitution across synaptic partners) (Feinberg et al., 2008; Gordon and Scott, 2009) where non-functional fragments of membrane tethered GFP (mCD4-GFP^1-10^ and mCD4-GFP^11^) were expressed in dfb neurons (23E10-Gal4) and OA-VPM3 neurons (24E06-Gal4). We found highly localized GFP signal in FB6 layers showing direct synaptic connectivity (Supplemental video 1: https://doi.org/10.5281/zenodo.14976495).

We next focused on the VPM3 outputs to the MB network as indicated by the EM data (Supplementary Table 1). Unlike the central complex whose cell types, populations and functions have only been described coarsely, MB cell types and their specific role in sleep are much more well characterized. Further for each of the cell types listed as VPM3 outputs and EM id we can target specific GAL4 and split-GAL4 lines. The VPM3 neurons project widely into the MB network with axonal projection in the γ Kenyon cells and significant posterior expression in the calyces around the KC dendrites. VPM3 also forms connections with two output neurons (MBON5 and MBON11). Unbiased activation-based screening has previously identified γm Kenyon cells and MBON5 as wake-promoting and suppress sleep when activated(Sitaraman et al., 2015a)

Both MBON05 (γ4>γ1γ2) and MBON11 (γ1pedc>α/β) based on light microscopic data have significant output directed to the dendrites of other MBONs within the MB lobes, potentially providing pathways for a multi-layered feedforward MBON network(Aso et al., 2014a, 2014b). In addition to KCs and MBONs, dopamine neurons (DANs) innervate sleep-regulating MB compartments suggesting that VPM3 modulates multiple interconnected nodes of the MB circuit (Figure 6B and C).

To specifically address if VPM3 mediated wakefulness requires MB, we performed a genetic/neural epistasis experiment (Figure 6C) in which VPM3 neurons were thermogenetically activated using dTRPA1 driven by VPM3-LexA and downstream targets (KCs, DANs and MBONs) targeted by GAL4s and split-GAL4s) were inhibited using Shibire^ts^. We measured sleep at 21°C (24 hours, 12hL:12hD) on day 1 followed by temperature shift during nighttime to 31°C (12h light, 21°C and 12h dark, 31°C) on day 2. We compared nighttime sleep on day 1 (Figure 6D and E) and day 2 (Figure 6F and G). Flies expressing dTRPA1 in VPM3 neurons were crossed to empty split-GAL4 (sp-GAL4) and empty GAL4 (pBD) as control and showed significant nighttime sleep suppression (Figure 6D, E, F and G).

VPM3 activation with simultaneous inhibition of specific wake-promoting DANs (PAM and PPL, Figure 6H) neurons, KCs (γ and α’β’, Figure 6I) and MBONs (MBON γ4>γ1γ2 and MBON- β’2mp, MBON-β’2mp_bilateral, MBON-γ5β’2a, Figure 6J) partially repressed VPM3 induced sleep suppression. However, inhibition of MBON 11(γ1pedc>α/β) and MBON 10 (β′1) which are also downstream of VPM3 based on EM data, did not rescue nighttime sleep suppression induced by OA-VPM activation. Taken together, VPM3 mediates wakefulness by directly or indirectly activity wake promoting KCs, DANs and specific MBONs.

In addition to selecting specific driver lines for downstream neurons based on EM data we also used a different approach to find sleep-regulating neurons downstream of OA-VPM3. A previous study aimed at identifying octopamine receptor expressing neurons involved in aggression generated 34 GAL4 driver lines containing molecularly defined cis-regulatory modules (CRMs) for four known Drosophila octopamine receptor genes(Watanabe et al., 2017). Interestingly one of the GAL4 lines we used to target PAM neurons (R48B04) in Figure 6 contains OAMB cis regulatory module and was developed as part of the 34 GAL4 collection targeting octopamine receptor neurons.

Of the 33 GAL4 drivers we selected 25 that had somewhat sparse expression and minimal VNC expression. We first screened these 25 GAL4 lines (Figure 6_supplement 1 A and B) with dTRPA1 for sleep phenotypes at 21°C and 29°C. Total sleep at 21°C was not significantly different between tested genotypes but several lines had increases and decreased sleep duration at 29°C as compared to genotypic controls (Figure 6_supplement 1 A and B). We found three octopamine receptor GAL4 lines that increased sleep when activated and 7 driver lines that suppressed sleep. We selected eight driver lines that were wake promoting and had broad expression in neuropils innervated by OA VPM3 neurons (Figure 6_supplement 1 C-J) and performed cellular epistasis experiments described above.

Nighttime sleep suppression induced by VPM3 activation was inhibited by three octopamine receptor GAL4 lines (R21B06, R50A06 and R19G08 Figure 6_supplement 1 K and L). R21B06 and R50A06 have localized expressions in Kenyon cells, while R19G08 has expression in the fan shaped body layers. Taken together, the unbiased screening of octopamine receptor GAL4 lines and cellular epistasis experiments provides converging evidence to support a role for KCs and fan shaped body neurons in VPM3 mediated sleep. However, these lines specifically R19G08 and R50A06 had widespread expression beyond KCs and dfb neurons and were not used for subsequent activation-based experiments.

### Role of specific octopamine receptors in sleep regulation

We next turned our attention to specific octopamine receptors involved in sleep regulation. Given that activating of neurons labeled by CRM derived from the Oamb, Octβ1R and Octβ2R genes changes sleep duration, we asked if these receptors regulate sleep. We used two independent approaches: screening of nulls/hypomorphs of different octopamine receptors and pan-neuronal knockdown using gene specific RNAi. In both approaches, we used nulls/hypomorphs and RNAi constructs that have been validated for knockdown of receptors using qRT-PCR and behavioral experiments(Burke et al., 2012; Deshpande et al., 2022; El-Kholy et al., 2022; Han et al., 1998; Lee et al., 2009, 2003; Lim et al., 2014; Pang et al., 2022; Sujkowski et al., 2017; Yoshinari et al., 2020; Zhao et al., 2021; Zhou et al., 2012).

We screened nulls/hypomorphs that were outcrossed for six generations into the w1118 background. We found Oamb, Ocβ1R and Octβ2R mutants slept more as compared to control flies (Figure 7A and B). We also measured P(doze), the probability that an active fly will go into quiescent mode and found it to be significantly higher in one of the OAMB mutants and Octβ2R mutant suggesting increased sleep pressure. Further for both lines, P(wake) is lower compared to controls and other mutants (Octβ1R and Octβ3R) suggesting increased sleep depth (Figure 7C and D).

**Figure 7:**
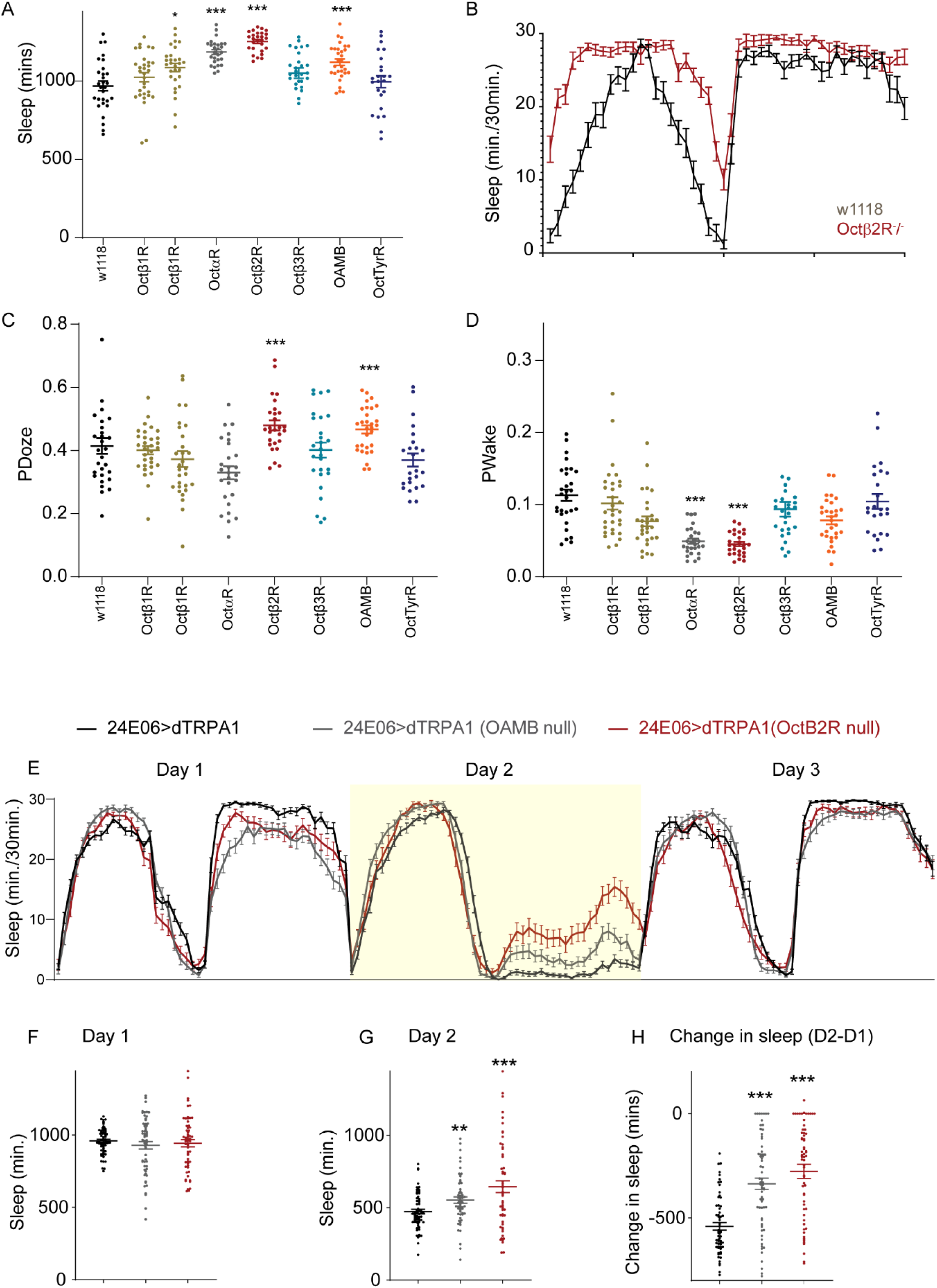
Specific octopamine receptors are involved in sleep regulation and block OA-VPM induced sleep loss. A. Sleep duration of null/hypomorphs of octopamine receptors ((Oct/tyrR(purple), Oamb (orange), Octα2R (grey) Octβ1R (green), Octβ2R (red) and Octβ3R(blue)) are shown. w1118 (black) is used as a genotypic control and data represents mean and sem and comparisons are made by Kruskal-Wallis test followed by Dunns multiple comparisons test. Statistical significances are indicated as ∗p < 0.05; ∗∗p < 0.01; ∗∗∗p < 0.001; ns, not significant. For each genotype no data was excluded, and data collected represents 2 or more independent trials. B. Representative sleep profile (mean and sem) of Octβ2R^-^/^-^ (red) and w1118 (black) flies. C and D. Pwake and PDoze of null/hypomorphs of 5 octopamine receptors ((Oct/Tyr(purple), Oamb(orange), Octα2R (grey) Octβ1R (green), Octβ2R (red) and Octβ3R(blue)) are shown. w1118 (black) is used as a genotypic control and data represents mean and sem and comparisons are made by Kruskal-Wallis test followed by Dunns multiple comparisons test. Statistical significances are indicated as ∗p < 0.05; ∗∗p < 0.01; ∗∗∗p < 0.001; ns, not significant. For each genotype no data was excluded, and data collected represents 3 or more independent trials. E. Sleep profile of flies expressing UAS-dTRPA1 in OA-VPM neurons (24E06-GAL4) in Octβ2R and OAMB heterozygous null background (day 1: 21, day 2: 29 (activation) and day 3: 21). F and G. Sleep duration on Day 1 (21) and Day 2 (29) of 24E06-GAL4>UAS-dTRPA1 (black, n=64), 24E06-GAL4>UAS-dTRPA1 in OAMB heterozygous null (grey, n=59) and Octβ2R background (red, n=57). H. Change in sleep induced by activation of OA-VPM neurons (day2-day1). Statistical significances are indicated as ∗p < 0.05; ∗∗p < 0.01; ∗∗∗p < 0.001; ns, not significant.

**Figure 7_Supplement 1:**
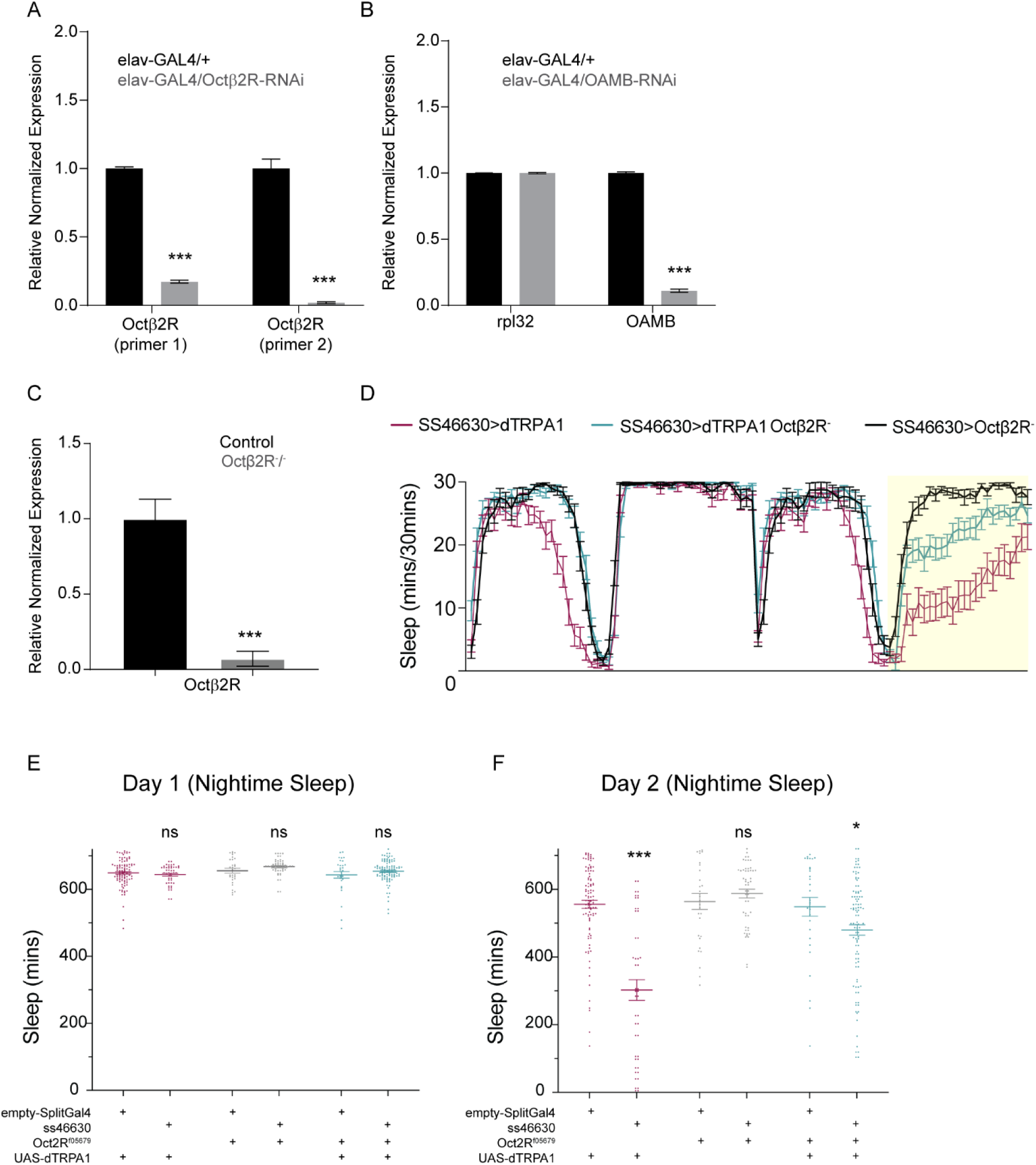
Octb2R receptor is required for sleep suppression induced by OA-VPM3 activation. A and B. Quantitative RT–PCR of Octβ2R and OAMB in flies with RNAi mediated pan-neuronal knockdown. Samples were compared using Mann Whitney U test and ∗∗∗p < 0.001. C. Quantitative RT–PCR of Octβ2R mutant and controls. Samples were compared using Mann Whitney U test and ∗∗∗p < 0.001. D. Representative sleep profile (mean and sem) of OA-VPM3>dTRPA1 flies in Octβ2R null and wild type background. The data represents baseline day 1 and day at 21^ο^C (daytime and nighttime) and day 2 (daytime at 21 ^ο^C and nighttime at 29 ^ο^C). E and F. Sleep profile of flies (SS46630>dTRPA1, SS46630> Octβ2R- and SS46630>dTRPA1, Octβ2R-). Day 1 and Day 2 represent daytime and nighttime sleep at 21 and 29^ο^C respectively and Day 1 represents daytime and nighttime sleep at 21 and Day 2 represents daytime at 21 and nighttime at 29 degrees respectively. Groups were compared made by Kruskal-Wallis test followed by Dunns multiple comparisons test. Statistical significances are indicated as ∗p < 0.05; ∗∗p < 0.01; ∗∗∗p < 0.001; ns, not significant.

Given the general role of octopamine neurons in arousal, it’s not surprising that several of the octopamine receptors are required to maintain wakefulness. To specifically address the role of these receptors in OA-VPM mediated arousal, we activated these neurons in OAMB and Octβ2R null mutant background and found that sleep loss is suppressed in these backgrounds (Figure 7E-H). We validated the efficacy of the mutants and RNAi knockdown by qPCR (Figure 7_supplement 1 A-C) and confirm that specific OA-VPM activation induced sleep suppression is rescued in Octβ2R mutant background (Figure 7_supplement 1 D-F).

In a complementary approach we used pan-neuronal knockdown of validated octopamine receptors RNAi lines targeted by elav-GAL4 to test a role for OAMB, Octβ1R, and Octβ2R in sleep regulation (Figure 8 A and B). Based on these results we targeted the RNAi lines that increased sleep pan-neuronally to more specific subsets within the MB (KCs, DANs and ONs).

**Figure 8:**
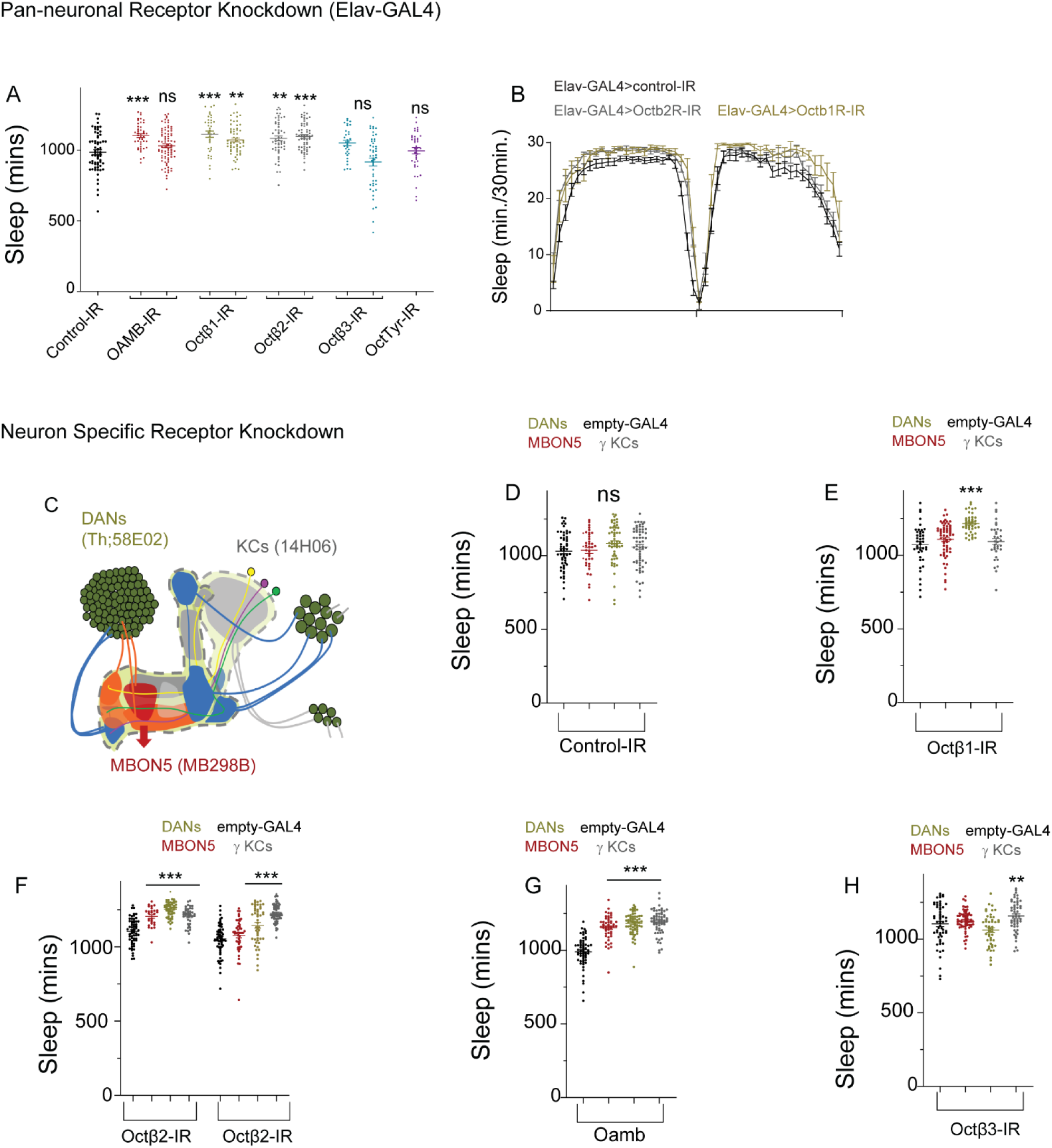
Targeted knockdown of specific octopamine receptor within MB cell types downstream of OA-VPM3 neurons increases sleep duration. A. Sleep duration of flies expressing elav-GAL4 and octopamine receptor RNAi lines. Oct/tyrR (purple,28332(purple, n=63), Oamb (red, 31233 (n=43) and 31171(n=82)) Octβ1R (green, 58179 (n=37) and 50701(n=58)), Octβ2R (grey, 50580 (n=54) and 34673 (n=63)) and Octβ3R(blue, 62283 (n=31) and 31108(n=48)) are shown. Elav-GAL4>36303 (black, empty RNAi) is used as a genotypic control and data is represented as scatter plots and comparisons are made by Kruskal-Wallis test followed by Dunns multiple comparisons test. Statistical significances are indicated as ∗p < 0.05; ∗∗p < 0.01; ∗∗∗p < 0.001; ns, not significant. For each genotype no data was excluded, and data collected represents 2 or more independent trials. B. Representative sleep profile (mean and sem) of Elav-GAL4>36303 (black), Elav-GAL4>UAS-Octβ2R-IR (red) and Elav-GAL4>UAS-Octβ1R-IR (green) C. The schematic of mushroom body where Kenyon cell (KC) bodies are represented grey and blue circles depending on the projection lobe. KCs dendrites extend within the calyx and axons project to form α/β, α’/β’, and γ lobes (targeted by 14H06-GAL4). PAM and PPL1 dopaminergic neurons (DANs) extensively innervate the lobes (green, Th;58E02). Mushroom body output neurons (MBON5), a key OA-VPM3 output is shown in red (MB298B). D-H. Sleep duration of flies expressing split-pBD (empty-GAL4, black), MB298B (MBON5, red), Th;58E02 GAL4 (DANs, Green), 14H06-GAL4 (γKCs, grey) and octopamine receptor RNAi lines (RNAi control (36303), Octβ1R (58179), Octβ2R (50580, 34673), Octβ3R (31108), and OAMB (31233)). Empty-GAL4 expressing specific receptor RNAi line is used as a genotypic control and data is represented as mean and sem and comparisons are made by Kruskal-Wallis test followed by Dunns multiple comparisons test. Statistical significances are indicated as ∗p < 0.05; ∗∗p < 0.01; ∗∗∗p < 0.001; ns, not significant. For each genotype no data was excluded, and data collected represents 2 or more independent trials. Number of flies for different genotypes is as follows: empty>36303 (n=54), MBON5>36303 (n=39), DAN>36303 (n=54) and γKCs>36303 (n=60). Empty>58179 (n=42), MBON5>58179 (n=63), DANs>58179(n=43) and γKCs>58179 (n=34). Empty>50580 (n=62), MBON5>50580(n=28), DANs>50580(n=49) and γKCs>50580 (n=39). Empty>34673 (n=62), MBON5>34673(n=46), DANs>34673 (n=49) and γKCs>34673 (n=64). Empty>31233 (n=58), MBON5>31233(n=50), DANs>31233 (n=63) and γKCs>31233 (n=62). Empty>31108 (n=58), MBON5>31108(n=63), DANs>31108 (n=47) and γKCs>31108 (n=52).

We used RNAi lines (OAMB/31233, Octβ1R/58179, Octβ2R/50580/34673 and Octβ3R/31108). that elevated sleep when expressed pan-neuronally and targeted them to the top output neuronal classes based on EM data and cellular epistasis experiments we conducted. Specifically, we targeted KCs (14H06-GAL4 with strong expression in γ Kenyon cells), dopamine neurons (targeted by Th-GAL4;58E02-GAL4) and MBON 5 (γ4>γ1γ2, MB298B) and measured sleep. Knockdown of Octβ2R, Octβ3R and OAMB increased sleep duration in MBON5 as compared to controls suggesting the role of multiple octopamine receptors in these neurons. Like MBON 5, knockdown of Octβ2R, Octβ3R and OAMB in KCs increased sleep. However, knockdown of all 4 octopamine receptors Octβ1R, Octβ2R, Octβ3R and OAMB in dopamine neurons increased sleep duration.

Together these data suggest that multiple octopamine receptors function within different subsets of neurons downstream of OA-VPM3 in regulating wakefulness, given the highly interconnected nature of these neurons, it’s highly likely that octopamine release activates multiple different nodes of the MB and produce concomitant signaling and wakefulness.

### Central complex inputs regulate activity of OA-VPM3 neurons

Based on the data presented we propose a model where OA-VPM3 function as sex specific arousal promoting neurons that increase courtship drive and suppress sleep by activating multiple wake-promoting neurons of the mushroom body including γ Kenyon cells, MBON 5 and DANs. The EM hemibrain data also reveals that FB6A and FB6C are reciprocally connected with OA-VPM3 and are key inputs. The targeting tools used for dfb cell types have extensive VNC expression and are not cell type specific, such that the commonly used 23E10-Gal4 has at least 9 cell types (Hulse et al., 2021) that include several FB6 cell classes. Additionally, the vnc neurons in this GAL4 line itself are sleep promoting confounding the role of dfb neurons in tracking sleep need (Jones et al., 2023). These inconsistencies have led to re-examining the precise role of these neurons in sleep homeostasis (De et al., 2023). However, in dfb neurons ATP levels rise after waking and predispose them to heightened oxidative stress linking enforced wakefulness to oxidative stress and altered mitochondrial dynamics(Sarnataro et al., 2024). The body of evidence makes FB6 neurons upstream of OA-VPM3 a critical node to understand how sleep/wake/arousal signals are relayed.

To directly address if dfb neurons that include FB6 classes are inhibitory to OA-VPM3 neurons we activated dfb neurons targeted by 23E10-LexA expressing CsChrimson and recorded calcium transients from VPM3 neurons targeted by 24E06-Gal4 and expressing GCamp6s. We found that red light stimulation reduced basal calcium fluorescence in the VPM3 neurons (Figure 9B-D). We also confirmed the synaptic connection in fb layers between these lines using GRASP (GFP reconstitution across synaptic partners) (Figure 9A and supplementary video 1: https://doi.org/10.5281/zenodo.14976495).

**Figure 9:**
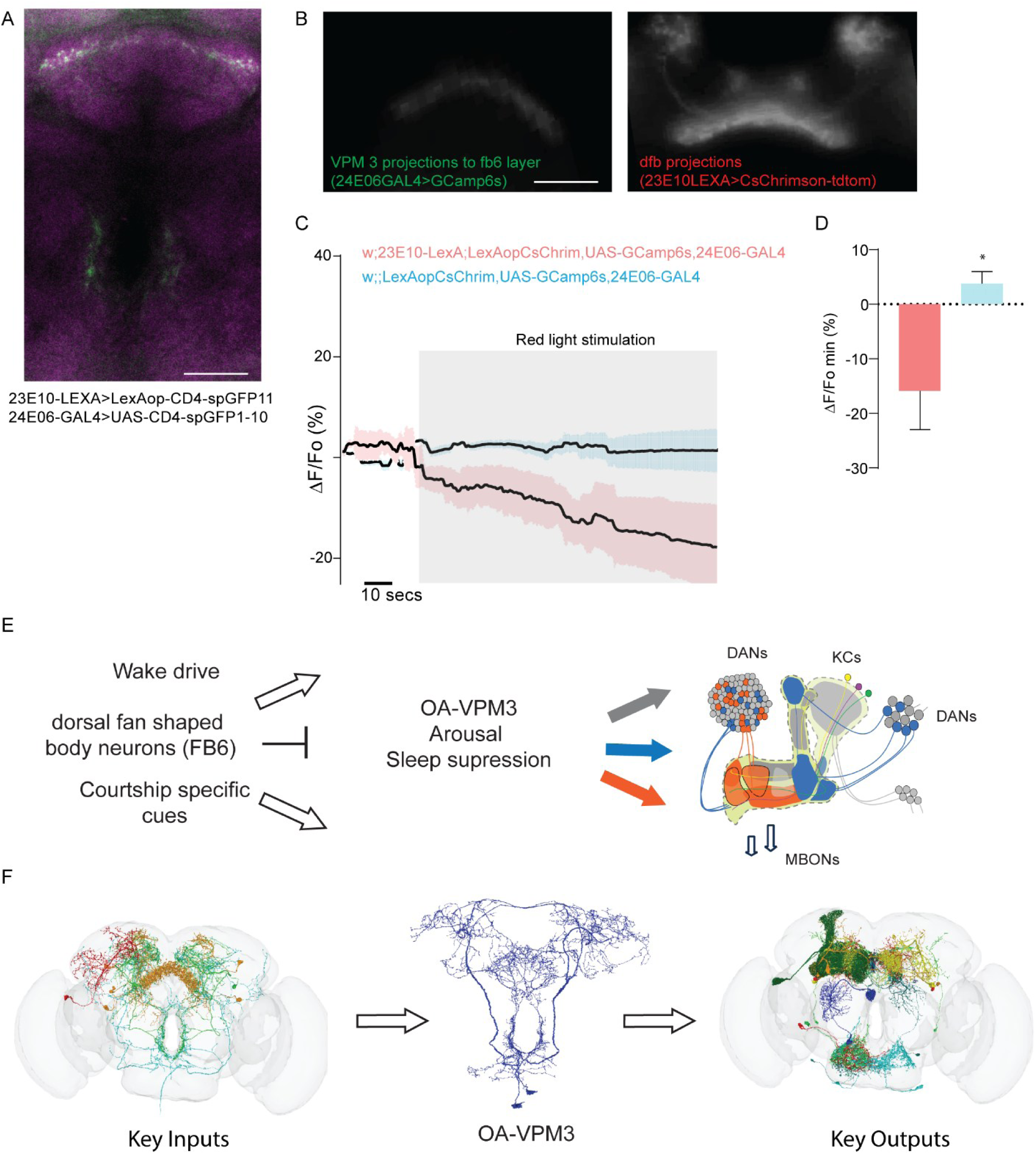
OA VPM3 neurons receive inhibitory input from fb6 neurons and connect key sleep regulating microcircuits between CX and MB. A. Central section of whole mount of male fly brain showing GRASP, Split-GFP reconstitution across synaptic partners-based connectivity between dorsal fan shaped body neurons targeted by 23E10-LexA and OA-VPM3 neurons targeted by 24E06-GAL4. Maximum intensity projection of the confocal image is shown, and scale bar represents 20um. B. Left and right panels showing GCamp6s expression in fb6 layer innervations of OA-VPM3 neurons and dfb (dorsal dan shaped body projections) of 23E10-LexA expressing CsChrimson-tdTomato. We used this region as a region of interest to measure changes in VPM3 activity as a function of dfb activation in male fly brains. C. Calcium transient (average traces, mean and sem) of normalized changes in GCamp6s fluorescence. Red light stimulation was provided 30 seconds after the start of recording to get a baseline fluorescence. Control group comprised of GCamp6s recording in flies with all transgenes except 23E10-LexA to drive CsChrimson. D. Bar graphs representing experimental (w,23E10-LexA; UAS-GCamp6s, CsChrimson-tdtomato,24E06-GAL4) and control (w,+;UAS-GCamp6s,CsChrimson-tdtomato,24E06-GAL4). Groups were compared using Mann Whitney U test (p=0.014, n=8 brains). E. Schematic of OA-VPM3’s role in mediating arousal and sleep suppression between central complex fan shaped body layers and mushroom body microcircuits implicated in sleep regulation. F. Flywire based filtering (10 or more synapses) of key inputs and outputs of OA-VPM3 neurons to illustrate compartmentalized input-output architecture of signals relayed via OA-VPM3 neurons.(flywire resource, Dorkenwald et al., 2024; Schlegel et al., 2024)

## Materials and Methods

### Flies

All Stocks and crosses were either reared at 21°C or 25°C and 50% humidity and maintained on a 12hr:12hr light: dark cycle in standard cornmeal media (https://bdsc.indiana.edu/information/recipes/bloomfood.html, Agar 0.56%, Cornmeal 6.71%, Inactivated Yeast 1.59%, Soy Flour 0.92%, Corn Syrup 7.00% per liter of food). Adult male and female Drosophila melanogaster of ages 4-10 days old were used for all experiments and group housed prior to experiments. All GMR GAL4, Split-GAL4, and LexA lines were obtained from the Bloomington Drosophila Stock Center or Janelia Research Campus. Flies used in experiments are listed in Table 2.

**Table 2:** List of fly lines used in the experiments:

| Reagent type<br>(species) or<br>resource | Genotype | Source |
| --- | --- | --- |
| Strain, strain<br>background<br>(*Drosophila*<br>*melanogaster*) | w[1118]; P{y[+t7.7]<br>w[+mC]=p65.AD.Uw}attP40; P{y[+t7.7]<br>w[+mC]=GAL4.DBD.Uw}attP2 | 79603 (BDSC) |
| Strain, strain<br>background<br>(*Drosophila*<br>*melanogaster*) | w[1118]; P{y[+t7.7]<br>w[+mC]=GAL4.1Uw}attP2 | 68384 (BDSC) |
| Strain, strain<br>background<br>(*Drosophila*<br>*melanogaster*) | MB021B (w; 24E06-p65ADZp in attP40;<br>95A10-ZpGdbd in attP2/TM6B) | 68297 (BDSC) |
| Strain, strain<br>background<br>(*Drosophila*<br>*melanogaster*) | MB022B (w; 24E06-p65ADZp in attP40;<br>TDC2-ZpGdbd in attP2/TM6B) | 68298(BDSC) |
| Strain, strain<br>background<br>(*Drosophila*<br>*melanogaster*) | ss46630 (w; 77F03-p65ADZp in attP40;<br>VT058566-ZpGdbd in attP2) | Dr. Gerry Rubin |
| Strain, strain<br>background<br>(*Drosophila melanogaster*) | ss46603( w; VT019391-p65ADZp in<br>attP40/CyOTb; VT058566-ZpGdbd in<br>attP2) | Dr. Gerry Rubin |
| Strain, strain<br>background<br>(*Drosophila melanogaster*) | w[1118]; P{y[+t7.7]<br>w[+mC]=GMR95A10-GAL4}attP2 | 41326 (BDSC) |
| Strain, strain<br>background<br>(*Drosophila melanogaster*) | w[1118]; P{y[+t7.7]<br>w[+mC]=GMR24E06-GAL4}attP2 | 48051 (BDSC) |
| Strain, strain<br>background<br>(*Drosophila melanogaster*) | w+; P{y[+t7.7] w[+mC]=UAS-<br>TrpA1(B).K}attP16 | Janelia Research<br>Campus |
| Strain, strain<br>background<br>(*Drosophila melanogaster*) | w;UAS-fruMIR (III) | Janelia Research<br>Campus (Bruce Baker<br>Lab) |
| Strain, strain<br>background<br>(*Drosophila melanogaster*) | w; P{10XUAS-IVS-mCD8::GFP}su(Hw)attP5 | 32188 (BDSC) |
| Strain, strain<br>background<br>(*Drosophila melanogaster*) | w+; P{w[+mC]=UAS-Hsap\KCNJ2.EGFP}7 | Janelia Research<br>Campus |
| Strain, strain<br>background<br>(*Drosophila melanogaster*) | w[*]; P{w[+mC]=LexAop-CD8-GFP-2A-CD8-GFP}2; P{w[+mC]=UAS-mLexA-VP16-NFAT}H2, P{w[+mC]=lexAop-rCD2-GFP}3/TM6B, Tb[1] | 66542 (BDSC) |
|  | w[1118] | 5905 (BDSC) |
| Strain, strain<br>background<br>(*Drosophila melanogaster*) | w[*]; P{w[+mC]=ple-GAL4.F}3 | 8848 (BDSC) |
| Strain, strain<br>background<br>(*Drosophila melanogaster*) | w[1118]; P{y[+t7.7]<br>w[+mC]=GMR48B04-GAL4}attP2 | 50347 (BDSC) |
| Strain, strain<br>background<br>(*Drosophila melanogaster*) | w[1118]; P{y[+t7.7]<br>w[+mC]=GMR58E02-GAL4}attP2 | 41347 (BDSC) |
| Strain, strain<br>background<br>(*Drosophila melanogaster*) | w[1118]; P{y[+t7.7]<br>w[+mC]=GMR15A04-GAL4}attP2 | 48671(BDSC) |
| Strain, strain<br>background<br>(*Drosophila melanogaster*) | w[1118]; P{w[+mC]=Ddc-GAL4.L}4.3D | 7010 (BDSC) |
| Strain, strain<br>background<br>(*Drosophila melanogaster*) | w[1118]; P{y[+t7.7]<br>w[+mC]=GMR19B03-GAL4}attP2 | 49830 (BDSC) |
| Strain, strain<br>background<br>(*Drosophila melanogaster*) | w[1118]; P{y[+t7.7]<br>w[+mC]=GMR35B12-GAL4}attP2 | 49822 (BDSC) |
| Strain, strain<br>background<br>(*Drosophila melanogaster*) | w[1118]; P{y[+t7.7]<br>w[+mC]=GMR14H06-GAL4}attP2 | 48667 (BDSC) |
| Strain, strain<br>background<br>(*Drosophila melanogaster*) | w[1118]; P{y[+t7.7] w[+mC]=R13F04-<br>GAL4.DBD}attP2 PBac{y[+mDint2]<br>w[+mC]=R93D10-p65.AD}VK00027<br>(MB112C) | 68263 (BDSC) |
| Strain, strain<br>background<br>(*Drosophila melanogaster*) | w [1118]; P{y[+t7.7] w[+mC]=R53H03-<br>GAL4.DBD}attP2 PBac{y[+mDint2]<br>w[+mC]=R30C01-p65.AD}VK00027<br>(MB078C) | Dr. Gerry Rubin |
| Strain, strain<br>background<br>(*Drosophila melanogaster*) | w[1118]; P{y[+t7.7] w[+mC]=R14C08-<br>p65.AD}attP40/CyO; P{y[+t7.7]<br>w[+mC]=R15B01-<br>GAL4.DBD}attP2/TM6B, Tb[1]<br>(MB011B) | 68294 (BDSC) |
| Strain, strain<br>background<br>(*Drosophila melanogaster*) | w[1118]; P{y[+t7.7] w[+mC]=R53C03-<br>p65.AD}attP40; P{y[+t7.7]<br>w[+mC]=R24E12-<br>GAL4.DBD}attP2/TM6B, Tb[1]<br>(MB298B) | 68309 (BDSC) |
| Strain, strain<br>background<br>(*Drosophila melanogaster*) | w[1118]; P{y[+t7.7]<br>w[+mC]=GMR19G08-GAL4}attP2 | 48863 (BDSC) |
| Strain, strain<br>background<br>(*Drosophila melanogaster*) | w[1118]; P{y[+t7.7]<br>w[+mC]=GMR49H01-GAL4}attP2 | 38711 (BDSC) |
| Strain, strain<br>background<br>(*Drosophila melanogaster*) | w[1118]; P{y[+t7.7]<br>w[+mC]=GMR50A06-GAL4}attP2 | 38722 (BDSC) |
| Strain, strain<br>background | w[1118]; P{y[+t7.7]<br>w[+mC]=GMR21B06-GAL4}attP2 | 49857 (BDSC) |
| *(Drosophila melanogaster)* |  |  |
| Strain, strain<br>background<br>*(Drosophila melanogaster)* | w[1118]; P{y[+t7.7]<br>w[+mC]=GMR47A04-GAL4}attP2 | 50286 (BDSC) |
| Strain, strain<br>background<br>*(Drosophila melanogaster)* | w[1118]; P{y[+t7.7]<br>w[+mC]=GMR47H03-GAL4}attP2 | 50331 (BDSC) |
| Strain, strain<br>background<br>*(Drosophila melanogaster)* | w[1118]; P{y[+t7.7]<br>w[+mC]=GMR47H08-GAL4}attP2 | 50335 (BDSC) |
| Strain, strain<br>background<br>*(Drosophila melanogaster)* | w[1118]; P{y[+t7.7]<br>w[+mC]=GMR48A07-GAL4}attP2 | 50340 (BDSC) |
| Strain, strain<br>background<br>*(Drosophila melanogaster)* | w[1118]; P{y[+t7.7]<br>w[+mC]=GMR48E03-GAL4}attP2 | 50368 (BDSC) |
| Strain, strain<br>background | w[1118]; P{y[+t7.7]<br>w[+mC]=GMR50A05-GAL4}attP2 | 38721 (BDSC) |
| *(Drosophila melanogaster)* |  |  |
| Strain, strain<br>background<br>*(Drosophila melanogaster)* | w[1118]; P{y[+t7.7]<br>w[+mC]=GMR19H07-GAL4}attP2 | 48867 (BDSC) |
| Strain, strain<br>background<br>*(Drosophila melanogaster)* | w[1118]; P{y[+t7.7]<br>w[+mC]=GMR20C11-GAL4}attP2 | 48887(BDSC) |
| Strain, strain<br>background<br>*(Drosophila melanogaster)* | w[1118]; P{y[+t7.7]<br>w[+mC]=GMR21A05-GAL4}attP2 | 48921 (BDSC) |
| Strain, strain<br>background<br>*(Drosophila melanogaster)* | w[1118]; P{y[+t7.7]<br>w[+mC]=GMR47C08-GAL4}attP2 | 50298 (BDSC) |
| Strain, strain<br>background<br>*(Drosophila melanogaster)* | w[1118]; P{y[+t7.7]<br>w[+mC]=GMR47D08-GAL4}attP2 | 50305 (BDSC) |
| Strain, strain<br>background | w[1118]; P{y[+t7.7]<br>w[+mC]=GMR49A02-GAL4}attP2 | 50398 (BDSC) |
| *(Drosophila melanogaster)* |  |  |
| Strain, strain<br>background<br>*(Drosophila melanogaster)* | w[1118]; P{y[+t7.7]<br>w[+mC]=GMR49C04-GAL4}attP2 | 50415 (BDSC) |
| Strain, strain<br>background<br>*(Drosophila melanogaster)* | w[1118]; P{y[+t7.7]<br>w[+mC]=GMR49G05-GAL4}attP2 | 38706 (BDSC) |
| Strain, strain<br>background<br>*(Drosophila melanogaster)* | w[1118]; P{y[+t7.7]<br>w[+mC]=GMR68B12-GAL4}attP2 | 39463 (BDSC) |
| Strain, strain<br>background<br>*(Drosophila melanogaster)* | w[1118]; P{y[+t7.7]<br>w[+mC]=GMR68C01-GAL4}attP2 | 39464 (BDSC) |
| Strain, strain<br>background<br>*(Drosophila melanogaster)* | w[1118]; P{y[+t7.7]<br>w[+mC]=GMR68C02-GAL4}attP2 | 39465 (BDSC) |
| Strain, strain<br>background | w[1118]; P{y[+t7.7]<br>w[+mC]=GMR68C08-GAL4}attP2 | 39468 (BDSC) |
| *(Drosophila melanogaster)* |  |  |
| Strain, strain<br>background<br>*(Drosophila melanogaster)* | w[1118]; P{y[+t7.7]<br>w[+mC]=GMR68C10-GAL4}attP2 | 39470 (BDSC) |
| Strain, strain<br>background<br>*(Drosophila melanogaster)* | w[1118]; P{y[+t7.7]<br>w[+mC]=GMR24E06-lexA}su(Hw)attP5 | Janelia Research<br>Campus |
| Strain, strain<br>background<br>*(Drosophila melanogaster)* | w[1118];<br>PBac{w[+mC]=WH}Octbeta1R[f02819]/T<br>M6B, Tb[1] | 18589 (BDSC) |
| Strain, strain<br>background<br>*(Drosophila melanogaster)* | w[*];<br>Tl{RFP[3xP3.cUa]=Tl}Octbeta1R[attP] | 84554 (BDSC) |
| Strain, strain<br>background<br>*(Drosophila melanogaster)* | w[1118];<br>PBac{w[+mC]=WH}Octalpha2R[f03483] | 18659 (BDSC) |
| Strain, strain<br>background | w[1118];<br>PBac{w[+mC]=WH}Octbeta2R[f05679]/T<br>M6B, Tb[1] | 18896 (BDSC) |
| *(Drosophila melanogaster)* |  |  |
| Strain, strain<br>background<br>*(Drosophila melanogaster)* | w[*];<br>Mi{y[+mDint2]=MIC}Octbeta3R[MI06217<br>] | 43050 (BDSC) |
| Strain, strain<br>background<br>*(Drosophila melanogaster)* | w[*];<br>Tl{RFP[3xP3.cUa]=Tl}Oamb[attP]/TM6B<br>, Tb[1] | 84555 (BDSC) |
| Strain, strain<br>background<br>*(Drosophila melanogaster)* | w[*];<br>Mi{y[+mDint2]=MIC}CG43102[MI04839] | 38558 (BDSC) |
| Strain, strain<br>background<br>*(Drosophila melanogaster)* | P{w[+mW.hs]=GawB}elav[C155];<br>P{w[+mC]=UAS-Dcr-2.D}2 | 25750 (BDSC) |
| Strain, strain<br>background<br>*(Drosophila melanogaster)* | y[1] v[1]; P{y[+t7.7]=CaryP}attP2 | 36303 (BDSC) |
| Strain, strain<br>background | y[1] v[1]; P{y[+t7.7]<br>v[+t1.8]=TRiP.HMJ22156}attP40 | 58179 (BDSC) |
| *(Drosophila melanogaster)* |  |  |
| Strain, strain<br>background<br>*(Drosophila melanogaster)* | y[1] v[1]; P{y[+t7.7]<br>v[+t1.8]=TRiP.GLC01702}attP40 | 50580 (BDSC) |
| Strain, strain<br>background<br>*(Drosophila melanogaster)* | y[1] sc[*] v[1] sev[21]; P{y[+t7.7]<br>v[+t1.8]=TRiP.HMS01151}attP2 | 34673 (BDSC) |
| Strain, strain<br>background<br>*(Drosophila melanogaster)* | y[1] v[1]; P{y[+t7.7]<br>v[+t1.8]=TRiP.JF01573}attP2/TM3, Sb[1] | 31108 (BDSC) |
| Strain, strain<br>background<br>*(Drosophila melanogaster)* | y[1] v[1]; P{y[+t7.7]<br>v[+t1.8]=TRiP.JF01732}attP2 | 31233 (BDSC) |
| Strain, strain<br>background<br>*(Drosophila melanogaster)* | y[1] v[1]; P{y[+t7.7]<br>v[+t1.8]=TRiP.JF02967}attP2 | 28332 (BDSC) |
| Strain, strain<br>background | y[1] v[1]; P{y[+t7.7]<br>v[+t1.8]=TRiP.HMJ23640}attP40 | 62283 (BDSC) |
| *(Drosophila melanogaster)* |  |  |
| Strain, strain<br>background<br>*(Drosophila melanogaster)* | w[1118]; P{y[+t7.7]<br>w[+mC]=GMR23E10-lexA}attP40 | 52693 (BDSC) |
| Strain, strain<br>background<br>*(Drosophila melanogaster)* | w[1118]; P{y[+t7.7]<br>w[+mC]=13XLexAop2-IVS-<br>CsChrimson.mVenus}attP2 | 55139 (BDSC) |
| Strain, strain<br>background<br>*(Drosophila melanogaster)* | w[1118]; PBac{y[+mDint2]<br>w[+mC]=20XUAS-IVS-<br>GCaMP6s}VK00005 | 42749 (BDSC) |

### Behavioral Experiments

Details for each of the four different assays performed in this paper are listed below. In general, all experiments were conducted in incubators maintained at specified temperature and humidity conditions.

### Sleep experiments

Locomotor activity and sleep were recorded with the Drosophila Activity Monitor (DAM2) system from TriKinetics (Waltham, MA, USA). Briefly, 3- to 8-day-old male flies were individually transferred to narrow glass tubes (65 mm X 5 mm) containing 5% sucrose and 1.5% agarose, loaded into the DAM systems, allowed to acclimatize to activity monitors and food for at least 12 hours, and monitored for 3-7 days in a 12:12 LD cycle. Data was collected in 1-minute bins. Total sleep duration, Pwake, PDoze and activity was extracted from the locomotor data as described in(Donelson et al., 2012; Sitaraman et al., 2024; Vecsey et al., 2024). Sleep profiles were generated depicting average sleep (minutes per 30 min) for the days of the experiment and maintained in the same tube. For experiments involving neural stimulation temperature changes are described in the figures.

### Male courtship assay

Four to eight-day-old wild-type CS group housed and mated females were loaded individually into round two-layer chambers (diameter: 1 cm; height: 2.5 mm per layer) as courtship targets, and 4–8-day-old tester males (group housed and mated) were then gently aspirated into the chambers. Recording started after flies were loaded for 30 mins using a camcorder (Cannon VIXIA HF R800). Courtship tests were conducted at 29°C and 21°C with genotypic controls. Courtship index, which is the percentage of observation time a male fly performs courtship, was used to measure courtship to female targets, and measured manually and scoring was performed single blind. We focused on three different courtship measures following, wing extension and attempted copulation as described in (Chen et al., 2017).

### Female receptivity assay

Tester females of desired genotypes (4–8 days old, group housed and mated) were aspirated into round two-layer chambers (diameter: 1 cm; height: 2.5 mm per layer) and separated from 4- to 8-day-old wild-type CS males (group housed and mated) until courtship test for 15 mins. Recording started when the barrier was removed and continued for 30 mins using a camcorder (Cannon VIXIA HF R800). Courtship tests were conducted at 29°C and 21°C with genotypic controls. Female receptivity, which is the percentage of observation time a female fly is receptive to male directed courtship, was used to measure courtship to male targets, and measured manually and scoring was performed single blind. We focused on three different female measures pausing, ovipositor extrusion and copulation as described in(Chen et al., 2017; Yang et al., 2023a)

### Single fly assay

Single male flies of desired genotypes (4–8 days old, group housed and mated) were aspirated into round two-layer chambers (diameter: 1 cm; height: 2.5 mm per layer). Recording started after 15 mins to allow fly to acclimatize to the chamber environment and continued for 30 mins using a camcorder (Cannon VIXIA HF R800). Activity was measured by tracking the flies in each chamber using the OpenCV module in the Python to analyze the 30-minute video and then output XY-coordinate and distance data. The analysis program is available freely on GitHub (GitHub - cgoina/pysolo-tools). Male courtship, female receptivity and single fly assays were all conducted between 2:00pm to 7:00pm.

### Immunohistochemistry

Brains or ventral nerve cords of 6–10-day old flies were dissected in cold PBS and fixed in 2% paraformaldehyde for 1 hour at room temperature. After fixation, samples were washed thrice with 0.05% PBST (PBS containing 0.05% Triton X-100) for 10 minutes at room temperature (RT), and then blocked in 5% normal goat serum (NGS) solution for 2 hours or overnight at 4°C. After blocking, samples were incubated with the primary antibody solution for two days at 4°C. The dilution ratios for the primary antibodies used in this study were 1:1000 for anti-GFP and 1:30 for nc82. Samples were washed in 0.05% PBST thrice in 10-minute blocks and incubated with the secondary antibody solution for two days at 4°C. Finally, the samples were washed three times with 0.05% PBST for 30 minutes at mounted using SlowFade™ Gold Antifade Mountant and imaged using Leica SP8 confocal microscope. The following antibodies were used: rabbit polyclonal anti-GFP (1:1000; Invitrogen), mouse anti-nc82 (1:50; Developmental Studies Hybridoma Bank, Univ. Iowa), and cross adsorbed secondary antibodies to IgG (H+L): goat Alexa Fluor 488 anti-rabbit (1:800; Invitrogen) and goat Alexa Fluor 568 (1:400; Invitrogen). Representative images illustrating the expression patterns of each driver were chosen from among 5-7 dissected samples.

### Confocal Imaging

Eight-bit images were acquired using a Leica TCS SP5 laser scanning confocal microscope with a 40×/1.3 numerical aperture (NA) or 20×/0.7 NA objective and a 1-3- μm z-step size. Maximum intensity z-projection images were generated in Fiji, a version of ImageJ software. For CaLexA experiment, we quantified fluorescence signal for a sum of slices within a manually drawn ROI. Number of slices summed, thickness of slices, and ROI dimensions were identical between comparison groups. Background fluorescence intensity adjacent to the ROI was measured and subtracted. For sleep deprivation experiments, flies were mechanically stimulated for 6 s per min (pulses were applied at random) for 16 hr. using a vortexer mounting plate and multi-tube vortexer (Trikinetics) and brains were dissected within 30 minutes of being removed from the deprivation set up.

#### Ex-vivo imaging of adult brains

Adult brains were dissected in adult hemolymph like (AHL) saline and placed in a perfusion chamber (PC-H chamber, Siskiyou Inc, OR) in AHL solution (5 mM HEPES pH 7.2, 70 mM NaCl, 20 mM KCl, 1.5 mM CaCl_2_, 20 mM MgCl_2_, 19 mM NahCO_3_, 5 mM trehalose, and 115 mM sucrose). A time series of fluorescence images was acquired using an Olympus BX51W microscope with U Plan Aprochromat 40X water immersion objective. GCamp6s was excited with a 470 nm LED light source (X-Cite turbo multiwavelength system) and images were acquired using ORCA FLASH 4.0 V2 digital CMOS camera. For stimulation of Chrimson, a 633nm LED line was used, while simultaneously acquiring GCaMP6s fluorescence images with a 470 nm LED to measure changes in Ca 2+. At least 5 independent brain preparations were used for all live imaging experiments and the exact number of cells imaged are indicated in the figures. Changes in fluorescence were calculated as ΔF/F = ((Ft − Fo)/Fo) where Fo is defined as the average background-subtracted baseline fluorescence for the 10 frames preceding red light stimulation. All images were processed and quantified using CellSens (Olympus Inc.) and Fiji (Image J).

#### RNA analysis

To test the efficiency of knocking down of Octβ2R RNAi and Oamb RNAi transgenes we used qRT PCR. Flies expressing UAS-RNAi pan-neuronally (Elav-Gal4) were collected and heads were removed using razor blades for RNA isolation. Three independent sets of 30 heads were ground using a pestle (Kimble® Disposable pellet pestle) and RNA was isolated using QIAGen RNeasy kit based on manufacturer instructions. 14μL of total RNA from each sample was reverse transcribed with the iScript cDNA Synthesis Kits. Isolated RNA was treated with RNase free DNase to eliminate genomic DNA contamination and reverse transcriptase was added to for a total sample amount of 20μL. cDNA was then diluted into equal concentration (300 ng/uL) to perform qPCR. Each biological replicate was assayed to identify the expression level of the targeted gene (Octβ2R and Oamb), as well as the internal standard endogenous reference genes (rpl32) using iQ SYBR Green Supermix. 1μL of each diluted cDNA reaction was used in 10 μL qPCR reaction and a master mix was made and 8μL transferred into wells of a 96-well plate. 2μL of working primers were then added into each well and mixed. qPCR thermal cycling and fluorescent data acquisition were performed with a BioRad Cfx96 system for 30s at 95°C followed by 40 cycles between 95°C (15s) and 60°C (30s). After the cycles, a melt curve analysis was performed from 65°C to 95°C with a 0.5°C increment in every 5s. Cq values were measured using CFX Maestro Software. A ΔΔCq method was used to process this data to calculate relative gene expression for the experiment.

Primers used (All primers were diluted to a final concentration of 500nM in reaction)

rpl32 forward 5ʹ-CCG CTT CAA GGG ACA GTA TC-3ʹ
rpl32 reverse 5ʹ-GAC AAT CTC CTT GCG CTT CT-3ʹ
DmOctβ2R forward 5ʹ-TCC TGT GGT ACA CAC TCT CCA-3ʹ
DmOctβ2R reverse 5ʹ-CCA CCA ATT GCA GAA CAG GC-3ʹ
DmOamb forward 5’-AGTCTAGAGCGGTTATACAGCCGACCTA
DmOamb reverse 5’-AAGAATTCGGGCGGAGTACAGGACATAA

### Data Analysis

Statistical analyses were performed with Prism 8 (GraphPad Inc). For comparisons of two groups of normally distributed data, Student’s t tests were performed. For multiple comparisons of normally distributed data, one-way or two-way ANOVAs followed by Tukey’s post-hoc tests were performed. For multiple comparisons of non-normally distributed data, Kruskal-Wallis test followed by Dunn’s post-hoc test was performed. The number of flies per genotype and p values are indicated in individual figures and figure legends.

## Discussion

Although sleep and wakefulness are measured as binary states at the behavioral level, animals exhibit various degrees and variations of arousal during wakefulness. Various behaviors including feeding, courting, escaping predators or unfavorable environments are key for survival and at odds with sleep phases. Theoretical and experimental findings show that sleep circuits exhibit emergent properties such as integration of signals across different time scales, outputs that depend on strength/duration of inputs and several self-sustaining feedback loops (Bhalla and Iyengar, 1999).

Here we focused our attention on a class of OA-VPM3 neurons in the central brain that receives and sends information to key sleep and arousal regulating loci in MB and CX. Thermogenetic activation of OA-VPM3 neurons targeted by a split-GAL4 suppresses sleep specifically at nighttime and the phenotype is stronger in males as compared to females. Waking as a behavioral state induced by VPM3 activation in males induces a strong sleep drive and on the subsequent day flies sleep more than controls. The role of OA-VPM3 neurons in sleep homeostasis is also supported by our findings that increased sleep post-deprivation is associated with decreased activity in sleep deprived flies as compared to sleep replete controls measured by CaLexA-mediated GFP expression levels.

Previous studies showed that the activation of broader OA neurons driven by Tdc2-GAL4 promotes arousal/wakefulness but suppresses subsequent sleep homeostasis (Seidner et al., 2015). However, single cell targeting tools such as ours show that subsets of OA neurons are not merely associated with arousal but regulate sleep and or encode sleep drive. This is not surprising as each of the OA neuronal subsets have distinct and extensive projection patterns and the likelihood that all Tdc2+ neurons are active at the same time is highly unlikely and misleading in interpreting the role of these complex systems.

In our pursuit to understand how subsets of OA neurons regulate arousal states, promote wakefulness and encode sleep drive we focused on VPM neurons in the central brain. OA central brain neurons tend to have connections with the most distinct cell types both in terms of morphology and neuropil location. Indeed, of the top 20 neurons in hemibrain, three (OA Vum7a, 6a and OA-VPM3) have the highest quantity of distinct cell types among their direct synaptic partners(Langstaff et al., 2024).

The two VPM3 cells are extensively connected with MB and CX and regulate wakefulness by direct and indirect connectivity with γ-lobe and α′β′-lobes. Furthermore, they are functionally and physically connected to PAM dopamine neurons and MBON5 and require OAM and Octβ2R receptors to regulate sleep. Previous work has shown that OA signaling via OAMB in the KCs is sufficient to rescue learning(Kim et al., 2013; Zhou et al., 2012 Burke et al., 2012; Huetteroth et al., 2015). OA signaling in α/β KCs is necessary for aversive olfactory learning, and OA signaling in projection neurons is involved in appetitive olfactory learning (Sabandal et al., 2020). In addition, Octβ2-R in the dopaminergic MB-MP1 PPL1 neurons is needed to modulate negative DA signals for sweet taste learning and suppression of Octβ2R in α′/β′ KCs impairs anesthesia-resistant memory (ARM)(Burke et al., 2012; Wu et al., 2013). Taken together, analyzing receptor function in specific sets of KCs, MB output neurons (MBONs), and DANs shows OA-VPM3 neurons form multi layered connectivity with MB and regulate sleep via multiple microcircuits and require both OAMB and Octβ2R. This likely provides a strategy to efficiently reset the larger MB network and provide redundancy to support states of wakefulness and allow efficient and complete transition between sleep to wake states.

In addition to the MB, OA-VPM3 has extensive innervation in the central complex and based on the adult female brain (FAFB) connectome, VPM3 neurons receive significant input from fb6 layers. Like with MBONs, some of these connections are reciprocal. The tangential neurons targeted by GAL4 lines with prominent expression in FB6 layers have been shown to track sleep need and function as an interconnected network with ER5 neurons controlling sleep-wake states (Donlea et al., 2014, 2011; Liu et al., 2016; Pimentel et al., 2016). The sleep promoting dFB neurons form reciprocal connections with wake-promoting dopaminergic neurons (DANs) potentially implementing a flip-flop circuit motif, that helps state switching without unstable intermediate states (Pimentel et al., 2016; Venner and Fuller, 2018). However, the targeting tools used for dfb cell types have extensive VNC expression and are not cell type specific, such that the commonly used 23E10-Gal4 has at least 9 cell types (Hulse et al., 2021). Additionally, the vnc neurons in this GAL4 line itself are sleep promoting confounding the role of dfb neurons in tracking sleep need (Jones et al., 2023). These inconsistencies have led to re-examining the precise role of these neurons in sleep homeostasis (De et al., 2023). More recent evidence points out a more complex role for these neurons in sleep. Several transcripts associated with ATP synthesis and mitochondrial respiration are up-regulated in dorsal fan-shaped body (dfb) neurons post sleep-deprivation and altering mitochondrial fission and fusion in dfb neurons affects sleep. The connection between aerobic metabolism, mitochondrial dynamics and sleep in dfb neurons suggests a more complex and nuanced function of sleep.

The connectome data sets of female brain show extensive reciprocal connectivity between OA- VPM3 and dfb neurons (specifically fb6A and 6D). Our GRASP experiments and calcium imaging shows these neurons are connected in male brains and activation of dfb neurons inhibits OA-VPM3 activity. Based on EASI-FISH experiments and RNA-seq data, fb6 neurons and subsets release AstC and both glutamate and acetylcholine (Wolff et al., 2024).

We recently published activation phenotypes of combinations of dFB cell types that represented different but largely overlapping subsets of layer 6 and 7 FB tangential cell types and have no detectable VNC expression and found that fb6 and 7 layers represent intermingled populations of sleep- and wake-promoting neurons (Wolff et al., 2024). At the EM level, many of these connect with OA-VPM3 neurons bi-directionally and warrant further investigation.

In addition to fb6, OA-VPM3 neurons are broadly connected with other cell types of CX that include PFGs, hΔK, hΔB, hΔH etc. although these connections are sparser based on connectome datasets. In our recent work using an unbiased optogenetic and thermogenetic screen we identified PFGs as a sleep-promoting cluster that makes reciprocal connections with fb6A neurons and is downstream of OA-VPM3 neurons and has a sex specific phenotype and activation of PFGs in female, but not male flies increase sleep (Hulse et al., 2021; Wolff et al., 2024).

The differential arousal needs of male and female flies could potentially influence sex specific sleep pattens and the underlying circuits. Octopamine circuits are largely sexually monomorphic, and subsets are FruM positive and known to regulate sex-specific arousal behaviors such as aggression, courtship, and, egg laying (Andrews et al., 2014; Certel et al., 2010; Deshpande et al., 2022; Dierick, 2008; Hoyer et al., 2008; Jia et al., 2021; Lee et al., 2003; Lim et al., 2014; Machado et al., 2017; Pang et al., 2022; Rohrscheib et al., 2015; Sherer et al., 2020; Stevenson et al., 2005; Yoshinari et al., 2020; Zhou et al., 2012, 2008).

Although there is no direct evidence of connectivity between OA-VPM3 neurons and Fruitless positive P1 neurons that promote courtship and suppress sleep, the extensive projections of VPM3 neurons in SEZ and LH (lateral horn) region suggests potential involvement in sensory processing of courtship relevant cues. Recent work shows that OA-VPM3 activation promotes sugar reinforced short term memory and inhibits long term memory, further artificial and odor (and learning) evoked activity helps the flies prioritize a newer experience over existing ones(Kapoor and Waddell, 2024). This work along with ours where activation induces a courtship arousal in courtship satiated flies suggests that like nor-epinephrine in mammals, OA gates sensory processing to enact motivated perceptual behaviors (Kapoor and Waddell, 2024; McBurney-Lin et al., 2019).

Acute OA-VPM3 activation potentially acts to improve sensitivity in early and intermediate sensory processing nodes to prioritize and enact motivational behaviors and long-term activity promotes states of wakefulness to support these arousal behaviors. Mechanistic insights into how they regulate sleep and arousal across timescales require further investigation. Taken together, the small number, discrete organization and identified connectivity patterns of OA- VPM3 neurons allows for an in-depth understanding of how male courtship drive and sleep need are encoded and communicated. Further, the extensive reciprocal connectivity and ability of these neurons to modulate sleep via MB and CX microcircuits suggests a potential mechanism by which these regions act together in enacting states of sleep and rest.

## Acknowledgements

We thank Dr. Gerry Rubin (Janelia Research Campus, HHMI) for providing split-GAL4/hemi-driver stocks and sharing brain and VNC images. The monoclonal nc82 antibody was obtained from the Developmental Studies Hybridoma Bank, created by the NICHD of the NIH and maintained at The University of Iowa, Department of Biology, Iowa City, IA 52242. Several stocks were obtained from Bloomington Drosophila Resource funded by NIH P40OD018537. We would also like to thank the members of the Sitaraman lab Steven Buchert, Veronica Ramirez, Bridget Fitzgerald, Roxanne Moghaddam and Suwei Lin for lively discussions and many suggestions. The research was supported by NIH 2R15GM125073-03 (to D.S.) and NSF CAREER IOS 2042873 (to D.S.).

## Notes

https://doi.org/10.5281/zenodo.14976495

### Competing Interest Statement

The authors have declared no competing interest.

